# A temporally controlled sequence of X-chromosome inactivation and reactivation defines female mouse *in vitro* germ cells with meiotic potential

**DOI:** 10.1101/2021.08.11.455976

**Authors:** Jacqueline Severino, Moritz Bauer, Tom Mattimoe, Niccolò Arecco, Luca Cozzuto, Patricia Lorden, Norio Hamada, Yoshiaki Nosaka, So Nagaoka, Holger Heyn, Katsuhiko Hayashi, Mitinori Saitou, Bernhard Payer

## Abstract

The early mammalian germ cell lineage is characterized by extensive epigenetic reprogramming, which is required for the maturation into functional eggs and sperm. In particular, the epigenome needs to be reset before parental marks can be established and then transmitted to the next generation. In the female germ line, reactivation of the inactive X-chromosome is one of the most prominent epigenetic reprogramming events, and despite its scale involving an entire chromosome affecting hundreds of genes, very little is known about its kinetics and biological function.

Here we investigate X-chromosome inactivation and reactivation dynamics by employing a tailor-made *in vitro* system to visualize the X-status during differentiation of primordial germ cell-like cells (PGCLCs) from female mouse embryonic stem cells (ESCs). We find that the degree of X-inactivation in PGCLCs is moderate when compared to somatic cells and characterized by a large number of genes escaping full inactivation. Nevertheless, PGCLCs that fail to undergo X-inactivation show an abnormal gene expression signature and deficiencies in meiotic entry. Subsequent to X-inactivation we observe gradual step-wise X-reactivation, which is mostly completed by the end of meiotic prophase I. Cells deviating from these progressive kinetics and undergoing X-reactivation too rapidly fail to enter a meiotic trajectory. Our data reveals that a fine-tuned X-inactivation and -reactivation cycle is a critical feature of female germ cell developmental competence towards meiosis and oogenesis

## Introduction

The germ cell lineage is unique in its critical function to transmit genetic and epigenetic information from one generation to the next. In mice, primordial germ cells (PGCs), the precursors of eggs and sperm, are specified during early postimplantation development from somatic precursors in the proximal epiblast by inductive signals (Ohinata *et al*, 2009, 2005; Lawson *et al*, 1999). Thereafter, PGCs migrate and enter the future gonads where they receive sex-specific somatic signals, which determine the germ cell sex and promote differentiation towards a spermatogenic or oogenic fate (Spiller *et al*, 2017; Miyauchi *et al*, 2017). While in males, germ cells enter mitotic arrest and differentiate into prospermatogonia, in females, germ cells instead progress into meiosis and oogenesis.

A hallmark feature of early germ cell development is the extensive epigenetic reprogramming (Kurimoto & Saitou, 2019), characterized by global changes in histone marks (Hajkova *et al*, 2008; Seki *et al*, 2005), DNA demethylation and erasure of genomic imprints (Seisenberger *et al*, 2012; Shirane *et al*, 2016; Hajkova *et al*, 2002). This establishes an epigenetic naive state (Ohta *et al*, 2017), which is required in order for PGCs to progress towards gonadal germ cell fate (Hill *et al*, 2018) and to control their timing to enter female meiosis (Yokobayashi *et al*, 2013). Ultimately, this erasure of parental information allows the reestablishment of new paternal and maternal marks during spermatogenesis and oogenesis, respectively, which are critical for the competence of egg and sperm to facilitate embryonic development in the next generation (Ohta *et al*, 2017; Reik & Surani, 2015).

In addition to these global changes, another important epigenetic reprogramming event takes place in the female germline; the reversal of silencing of the inactive X chromosome by X-chromosome reactivation. While X-chromosome inactivation (Lyon, 1961; Galupa & Heard, 2018; Payer & Lee, 2008) is the process by which female mammals (XX) achieve X-linked gene dosage parity with males (XY), X-reactivation takes place specifically in pluripotent epiblast cells of the mouse blastocyst (Mak *et al*, 2004; Borensztein *et al*, 2017) and in PGCs during their migration and upon their entry into the gonads (Sugimoto & Abe, 2007; Chuva de Sousa Lopes *et al*, 2008). Therefore, while X-inactivation is associated with pluripotency exit and the differentiated state (Schulz *et al*, 2014), X-reactivation is a key feature of naive pluripotency and germ cell development (Payer, 2016; Talon *et al*, 2019; Pasque *et al*, 2014; Bauer *et al*, 2021; Janiszewski *et al*, 2019). X-reactivation in mouse PGCs is a multistep process, which initiates during PGC migration with downregulation of Xist, the long non-coding master regulator RNA of X-inactivation, and concomitant loss of the associated histone H3K27me3 mark from the inactive X (Sugimoto & Abe, 2007; Chuva de Sousa Lopes *et al*, 2008). This process is regulated by repression of the *Xist* gene by the germ cell transcription factor PRDM14 (Mallol *et al*, 2019; Payer *et al*, 2013) and potentially by other members of the pluripotency network such as NANOG or OCT4 (Navarro *et al*, 2008), which are all expressed during PGC development. Subsequently, X-linked genes become progressively reactivated during migration, with the process being completed after PGCs have reached the gonads, and following the initiation of oogenesis and meiosis (Sangrithi *et al*, 2017; Sugimoto & Abe, 2007). X-linked gene reactivation is thereby thought to be enhanced by a female-specific signal from gonadal somatic cells (Chuva de Sousa Lopes *et al*, 2008). Although the molecular nature of the X-reactivation-promoting signal is currently unknown, the timing of X-linked gene reactivation around meiotic entry and the dependency of both processes on a female somatic signal, suggests a potential mechanistic link. Until now it has not been formally tested, if, and to which degree, the X-inactivation status might impact the meiotic and oogenic potential of germ cells. Furthermore, previous studies on the X-inactivation and - reactivation dynamics during mouse germ cell development have been limited to few individual genes (Sugimoto & Abe, 2007) or have not been allelically resolved and therefore been unable to discriminate between transcripts expressed from either one or two X chromosomes (Sangrithi *et al*, 2017). Therefore a comprehensive analysis of X-inactivation and -reactivating kinetics and its functional relation to germ cell developmental progression is necessary to gain mechanistic insight.

Based on *in vitro* germ cell differentiation from mouse embryonic stem cells (ESCs) (Nakaki *et al*, 2013; Hayashi *et al*, 2011, 2012), we developed an X-chromosome reporter system (XRep) to study the kinetics of X-inactivation and - reactivation during germ cell development. We thereby provide a high-resolution allelic analysis of X-chromosome dynamics and discovered that germ cells with high meiotic and oogenic competence are characterized by a moderate degree of X-inactivation and gradual X-reactivation kinetics. In contrast, germ cells that failed to undergo X-inactivation or which reactivated the X chromosome too rapidly displayed abnormal gene expression and differentiation characteristics. Thus, we found first evidence that a controlled sequence of X-inactivation followed by X-reactivation to be a characteristic hallmark of normal female germ cells. This suggests that both dosage control and epigenetic reprogramming of the X chromosome may be critical steps required for female germ cell’s developmental progression towards meiosis and oogenesis.

## Results

### XRep, a tailor-made system for tracing X-chromosome dynamics during in vitro germ cell development

In order to achieve a better understanding of the X-chromosome dynamics during mouse germ cell development, we created a tailor-made in vitro model system called XRep (Fig. 1A). XRep combines the following features. First, it is based on a hybrid female embryonic stem cell (ESC) line containing one *Mus musculus* (X^mus^) and one *Mus castaneus* (X^cas^) X chromosome (Lee & Lu, 1999; Ogawa *et al*, 2008), allowing allele-specific determination of gene expression. Moreover, this line was shown to be karyotypically highly stable (Lee & Lu, 1999; Bauer *et al*, 2021), therefore preventing X-loss, a crucial prerequisite for X-inactivation and -reactivation studies. Additionally, the cell line contains a *Tsix* truncation (TST) on X^mus^, forcing non-random X inactivation of the X^mus^ upon cell differentiation (Ogawa *et al*, 2008; Luikenhuis *et al*, 2001). This enabled us to study the X-inactivation and -reactivation dynamics specifically of the X^mus^ chromosome, while the X^cas^ would remain constitutively active. Second, primordial germ cell-like cells (PGCLCs) can be obtained highly efficiently from XRep cells by doxycycline-inducible overexpression of the germ cell fate specifier transcription factors BLIMP1 (also known as PRDM1), PRDM14 and TFAP2C (also known as AP2γ) (Nakaki *et al*, 2013), therefore bypassing the need for addition of cytokines. Last, the X-chromosome status of XRep cells can be traced by dual X-linked reporter genes placed in the *Hprt*-locus (Wu *et al*, 2014), a GFP reporter on X^mus^ (XGFP) and a tdTomato reporter on X^cas^ (XTomato). This allows us to isolate distinct populations of cells, harboring either two active X chromosomes (XGFP+/XTomato+) or one inactive and one active X (XGFP-/XTomato+), using fluorescence-activated cell sorting (FACS). This gives us a unique advantage over *in vivo* studies, as it enables us to test the importance of X-inactivation and -reactivation for germ cell development by isolating and further culturing cells of different X-inactivation states. Taken together, this tailor-made system allows us to assess X-chromosome dynamics and its importance for female mouse germ cell development *in vitro*.

**Figure 1.**
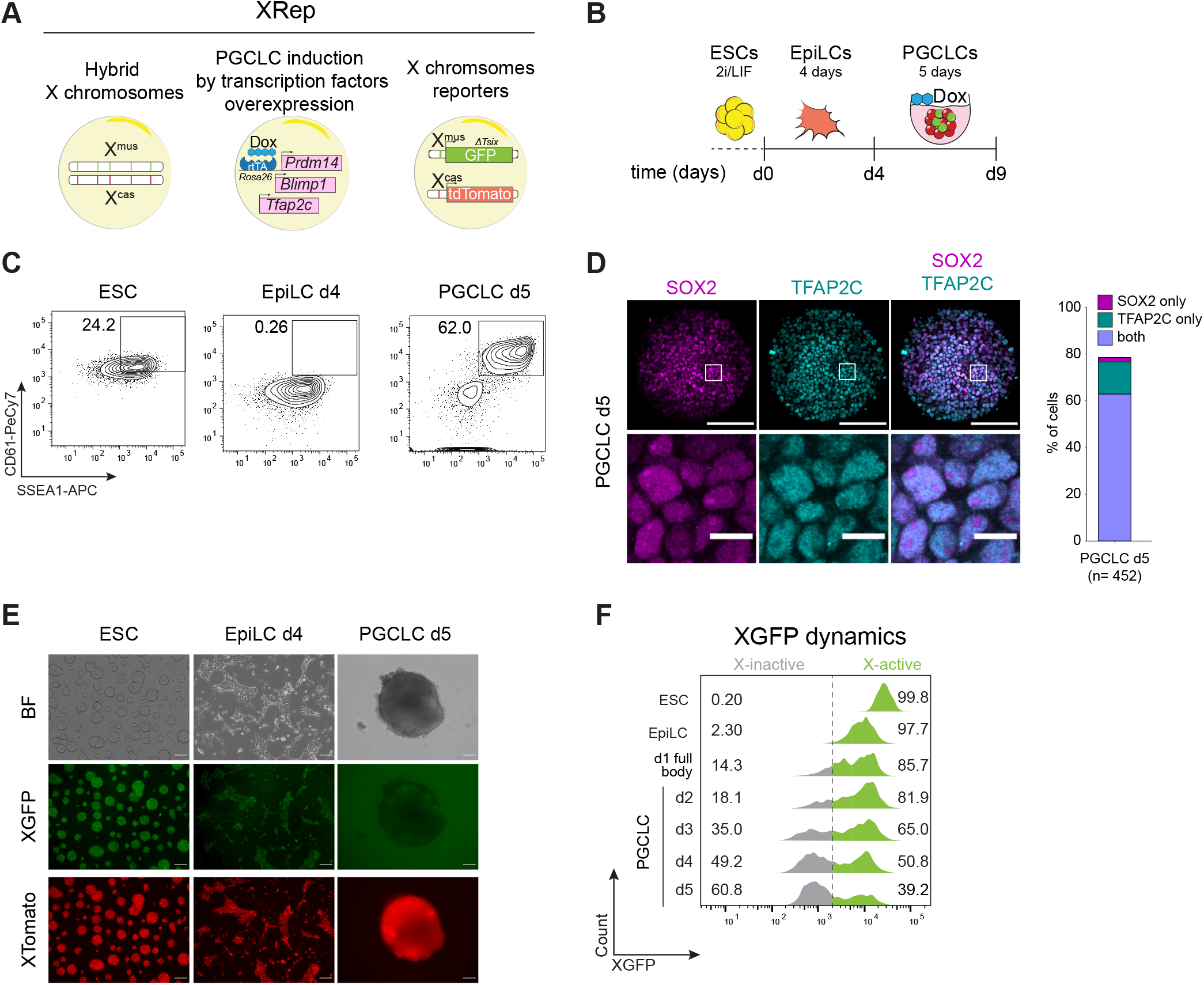
A tailor-made system to trace X-chromosome inactivation and reactivation dynamics during PGCLC induction. **(A)** Schematic representation of the features implemented in the XRep cell line. A hybrid background in which cells carry one X chromosome from M.m musculus (Xmus) and one from M.m castaneus (Xcas). The cell line carries an rtTA under the control of the Rosa26 locus and piggyBac transposon-based vectors with doxycycline (Dox)-responsive promoters driving the expression of Prdm14, Blimp1 and Tfap2c. The Xmus carries a GFP reporter and a truncation of the Tsix transcript while the Xcas carries a tdTomato reporter. **(B)** Overview of the adapted PGCLC differentiation timeline. Stages of the culture system are shown. **(C)** Representative FACS data of primordial germ cells specific surface markers CD61 and SSEA1 in ESCs, EpiLCs d4 and PGCLCs d5. Numbers indicate the percentages of SSEA1+/CD61+ gated cells over time. Shown are contour plots gated on live cells. **(D)** Immunostaining of PGCLCs d5 cryosections for SOX2 (magenta) and TFAP2C (cyan). Barplot indicates the quantification of SOX2+ cells, TFAP2C+ cells, and SOX2+/TFAP2C+. n=452 cells, from n=1 PGCLC aggregate. The white squares represent the position of the magnified region at the bottom. Scale bar, 50 µm and 10 µm for the magnified region. **(E)** Representative culture showing the X-activity reporter during PGCLC induction. Images for bright field (BF), XGFP, and XTomato were taken for ESCs, EpiLC d4 and PGCLC d5. Scale bar, 50 μm. **(F)** Representative FACS data showing XGFP distribution during PGCLC induction. Numbers indicate the percentage of gated cells according to the XGFP status (gray = X-inactive, green = X-active). Dashed line indicates the transition from X-active to X-inactive according to XGFP levels. XGFP percentages from PGCLC d2 to PGCLC d5 are calculated from SSEA1+/CD61+ PGCLCs. Shown are histograms gated on live cells.

We first set out to assess competence for PGCLC differentiation of our XRep cell line. We slightly adapted published protocols (Hayashi & Saitou, 2013; Nakaki *et al*, 2013), by differentiating ESCs into epiblast-like cells (EpiLCs) for four days, as differentiation for two days did not yield PGCLCs with our XRep cells likely due to their specific genetic background (Fig. EV1A), followed by induction into PGCLCs for five days (Fig. 1B). We quantified PGCLC induction efficiency by FACS analysis, using SSEA1 and CD61 double-positive staining to mark successfully induced PGCLCs (Fig. 1C). At PGCLC day 5, we found ∼60% of the cell population to be double-positive for SSEA1/CD61, indicating a very high PGCLC induction efficiency when compared to the cytokine-based protocol (Hayashi & Saitou, 2013) and in line with previous observations on transcription factor-based PGCLC induction (Nakaki *et al*, 2013). To further assess the quality of our PGCLCs, we stained cryosections of PGCLC bodies at day 5 of induction for SOX2 and TFAP2C, both germ-line expressed transcription factors. We observed that more than 50% of cells were double-positive for SOX2 and TFAP2C (Fig. 1D), confirming PGCLC cell identity. We next wanted to assess X-inactivation kinetics using our XGFP and XTomato reporters. As expected, XTomato stayed active throughout the differentiation (Fig. EV1B). In contrast, we observed downregulation of the XGFP reporter at day 2 of PGCLC differentiation, with the XGFP-population gradually increasing until day 5 (Fig. 1E and F). Nevertheless, even at day 5, up to 40% of PGCLCs remained XGFP+ in our system. Despite this, the large majority of EpiLCs showed H3K27me3 foci indicating initiation of X-inactivation (Fig. EV1C). As H3K27me3 accumulation on the Xi was shown to initiate prior to gene silencing (Zylicz *et al*, 2019), this suggests that XGFP+ PGCLCs failed to undergo X-linked gene silencing rather than originating from a subpopulation that did not initiate X-inactivation. In summary, using our tailor-made XRep cell line, we could show that X-inactivation occurs early during PGCLC differentiation. Additionally, our system enables the isolation of distinct PGCLC populations either having undergone X-inactivation or harboring two active X, suggesting that PGCLC specification can occur in the absence of X-inactivation as well.

**Figure EV1.**
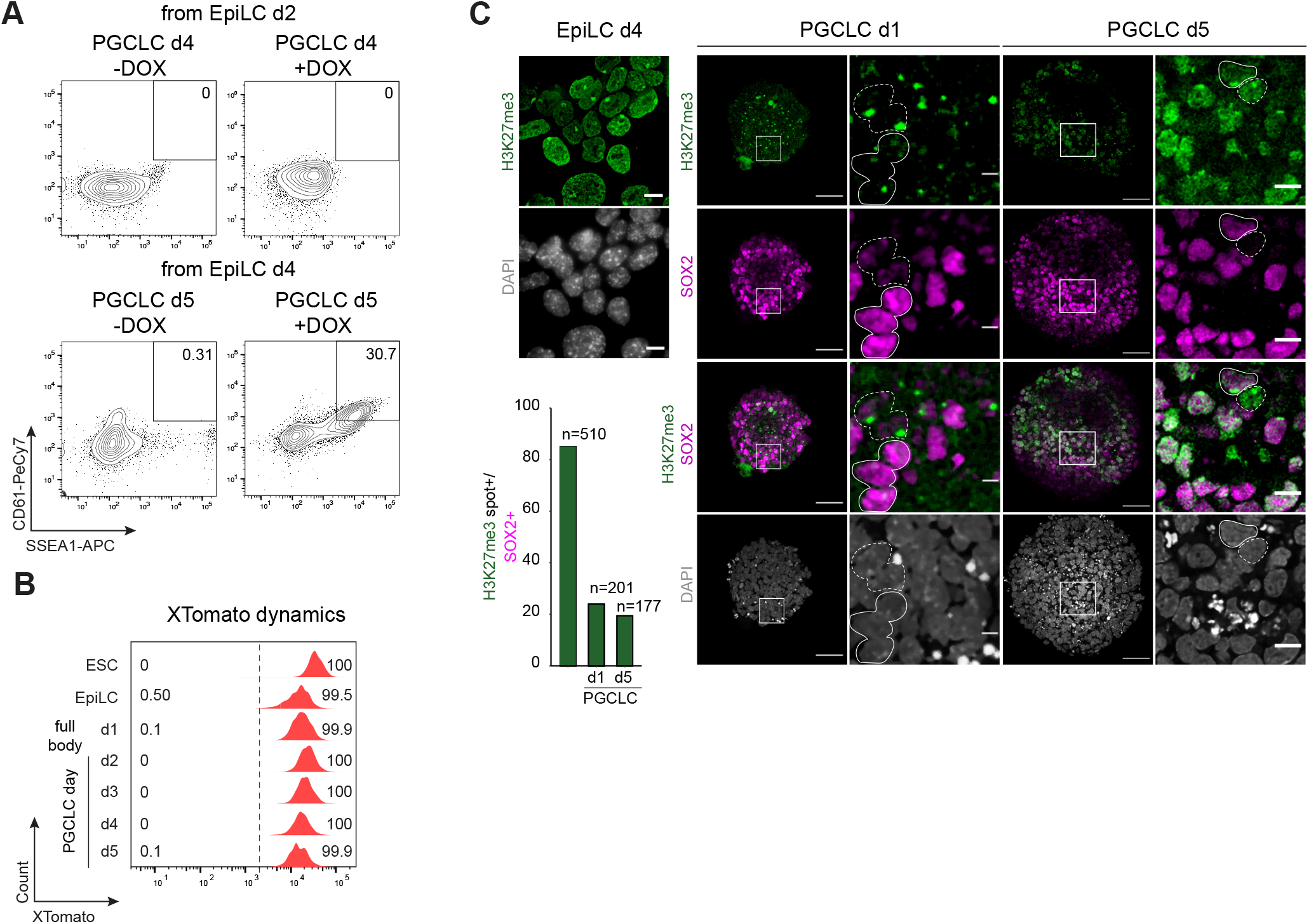
A tailor-made system to trace X-chromosome inactivation and reactivation dynamics during PGCLC induction. **(A)** Top panel shows representative contour plots of FACS analysis of PGCLC induction, without or with Dox, in PGCLC d4 induced from EpiLC d2. Bottom panel shows PGCLC d5 induced from EpiLC d4. The number indicates the gated germ cells identified by CD61 and SSEA1 signal. Shown are contour plots gated on live cells. **(B)** Representative FACS data showing XTomato distribution during PGCLC induction. Numbers indicate the percentage of gated cells according to the XTomato status (gray = X-inactive, red = X-active). Dashed line indicates the transition from X-active to X-inactive according to XTomato levels. XTomato percentages from PGCLC d2 to PGCLC d5 are calculated from SSEA1+/CD61+ PGCLCs. Shown are histograms gated on live cells. **(C)** Immunolabeling with antibodies against H3K27me3 (green) in EpiLCs, combined with SOX2 (magenta) in PGCLCs d1 and PGCLCs d5. Images show representative groups of cells showing H3K27me3 enrichment on the Xmus. Barplots indicate the percentage of cells having H3K27me3 accumulation. PGCLC d1 and PGCLC d5 H3K27me percentages are calculated from SOX2 positive cells. On top of the bars, the total cell number analysed from n=1 body is indicated. The white squares represent the position of the magnified region at the bottom. Dashed line indicates SOX2-/H3K27me3+ cells. Continuous line indicates SOX2+/H3K27me3+ cells. Scale bar, 50 µm and 10 µm for the magnified region.

### XGFP+ and XGFP-PGCLCs define distinct subpopulations

Having identified two distinct PGCLC populations, we set out to characterize the transcriptional changes taking place during differentiation. We induced EpiLCs from ESCs for 4 days and subsequently induced PGCLCs for 5 days, at which stage we isolated XGFP+ and XGFP-PGCLCs by FACS (Fig. EV2A). With these samples, we performed allele-specific RNA-sequencing on two biological replicates (different clones) with two technical replicates each. Principal component analysis (PCA) of the expression profiles showed a high coherence between replicates, with ESCs, EpiLCs and PGCLCs occupying distinct clusters (Fig. 2A). Moreover, we observed that XGFP+ and XGFP-PGCLCs clustered separately, indicating distinct expression profiles of the two populations. To exclude the possibility that the distinct clustering of PGCLC populations was influenced by the different X-status of the two, we repeated the PCA while eliminating X chromosome-linked genes from the analysis. We observed a highly similar clustering of samples with minimal changes in component variances (Fig. EV2B). In order to assess if transcriptional differences in XGFP+ and XGFP-PGCLCs could be explained by differences in developmental timing, we took advantage of published datasets of female *in vivo* PGCs from E9.5, E10.5, E11.5 and E12.5 embryos (Nagaoka *et al*, 2020) and compared expression profiles to our *in vitro* derived PGCLCs (Fig. 2B). PCA revealed a trajectory where PC1 defined the developmental timing of *in vivo* samples. We found that both PGCLC populations clustered around E10.5, with XGFP+ cells corresponding to a slightly advanced developmental stage. Therefore, as XGFP+ and XGFP-PGCLCs seemed to correspond to a similar developmental time point, we wanted to characterize their transcriptional differences in more detail. We performed differential gene expression analysis and could identify 2684 upregulated and 2437 downregulated genes in XGFP-PGCLCs, when compared to XGFP+ PGCLCs (Fig. 2C-E). Among the genes upregulated in XGFP-PGCLCs, we found early germ cell genes including *Blimp1 (Prdm1), Prdm14* and *Tfap2c*. In contrast, in XGFP+ PGCLCs we observed higher expression of pluripotency genes such as *Esrrb* and *Zfp42* and a subset of late germ cell genes like *Dazl*.

**Figure 2.**
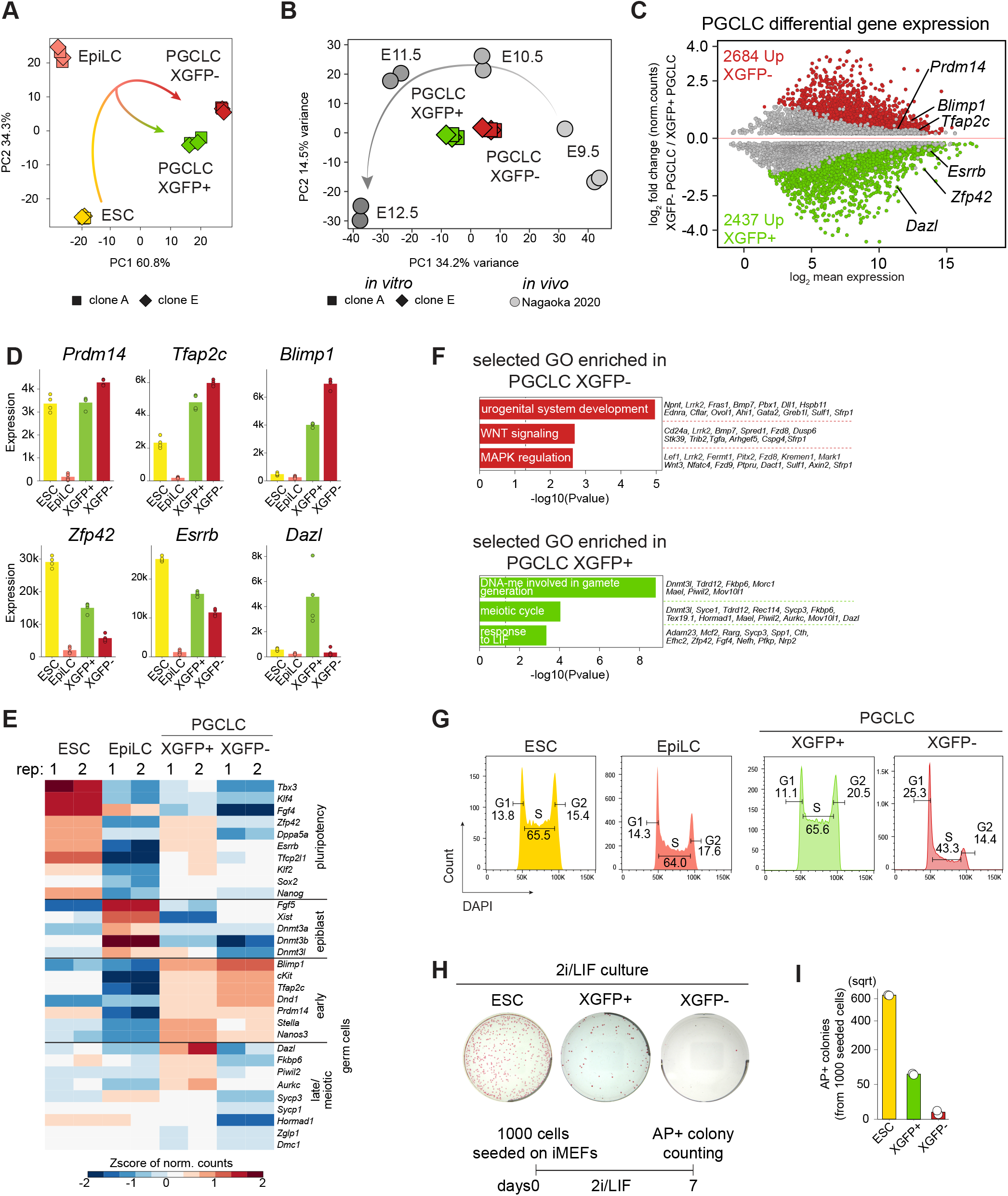
Gene expression analysis reveals two PGCLCs subpopulations. **(A)** PCA of gene expression dynamics during PGCLC differentiation. 4 biological replicates are shown. n = top 500 most variable genes. PGCLCs were sorted for SSEA1 and CD61 expression and further divided into XGFP+ and XGFP-. Axes indicate the variance. Arrows indicate hypothetical trajectory. Shapes indicate the biological clone (clone A11 = square, clone E9 = rhombus). **(B)** PCA of gene expression dynamics compared to in vivo samples from (Nagaoka et al, 2020). n = top 500 most variable genes, calculated including in vivo samples. Gray arrow indicates putative developmental trajectory. Shapes indicate the replicates (clone A11 = square, clone E9 = rhombus, in vivo samples = circle). **(C)** MA plot of differential gene expression changes between XGFP- and XGFP+ PGCLCs as determined by RNA-seq. Log2-mean expression (log2-normalized counts from DESeq2) on the x-axis and the log2-fold change on the y-axis are shown. Significantly upregulated and downregulated genes are highlighted in red and green respectively. False Discovery Rate (FDR) < 0.001. Non-significant genes with log2-mean expression between 0 and 0.2 were removed for easier plot visualization. **(D)** Expression levels (normalized DEseq2 counts) of selected differentially expressed genes between XGFP- and XGFP+ PGCLCs during the differentiation time course. Genes with FDR < 0.001 were considered significantly differentially expressed. Points indicate expression of individual biological replicates. **(E)** Heatmap of RNA-seq normalized counts showing the Zscore across PGCLC induction timepoints of 31 manually selected and manually ordered marker genes belonging to the categories reported on the side. **(F)** Selected GO terms enriched in XGFP-PGCLCs and XGFP+ PGCLCs. **(G)** FACS analysis of cell cycle using DAPI. Numbers indicate the percentage of cells in G1, S and G2/M respectively. **(H)** Alkaline phosphatase staining for ESC and XGFP+ PGCLC and XGFP-PGCLC grown for 7 days in 2i/LIF medium on immortalized feeder cells. **(I)** Barplot indicates the absolute numbers of Alkaline Phosphatase (AP) positive colonies in each cell type after 7 days of culture in 2i/LIF medium on immortalized feeder cells. Y-axis is in square root scale (sqrt) for easier plot visualization. Each white dot represents one technical replicate.

Moreover, when we performed functional annotation by gene ontology (GO) term analysis we observed enrichment for genes involved in urogenital system development, MAPK regulation and WNT signaling in XGFP-PGCLCs, while genes upregulated in XGFP+ PGCLCs were enriched for DNA methylation involved in gamete generation, meiotic cell cycle and response to LIF signaling (Fig. 2F). MAPK signaling is known to be inhibited by double X-dosage (Schulz *et al*, 2014; Song *et al*, 2019; Genolet *et al*, 2021), which might explain enrichment of this pathway in our XGFP-PGCLCs. LIF signaling on the other hand, which is enriched in our XGFP+ PGCLCs, is known to enable expression of the naive pluripotency network, which represses *Xist*, thereby promoting the active X state (Payer & Lee, 2014; Panda *et al*, 2020). Furthermore, enrichment for meiotic cell cycle genes in XGFP+ PGCLCs such as *Aurkc, Dazl* and *Piwil2*, suggests a premature activation of a subset of meiotic genes in XGFP+ PGCLCs. One characteristic feature of PGCLCs are changes in cell cycle progression and proliferation upon differentiation (Ohta *et al*, 2017), both of which are known to be affected by MAPK, as well as LIF signaling pathways (Onishi & Zandstra, 2015; Meloche & Pouysségur, 2007). We therefore performed cell cycle analysis using DAPI and found that ESCs, EpiLCs and XGFP+ PGCLCs shared highly similar profiles, with the majority of cells (>60%) residing in S phase. In contrast, XGFP-PGCLCs showed a decreased number of cells in S phase, concomitant with an increase of cells in G1, suggesting a slower proliferation of this population (Fig. 2G).

**Figure EV2.**
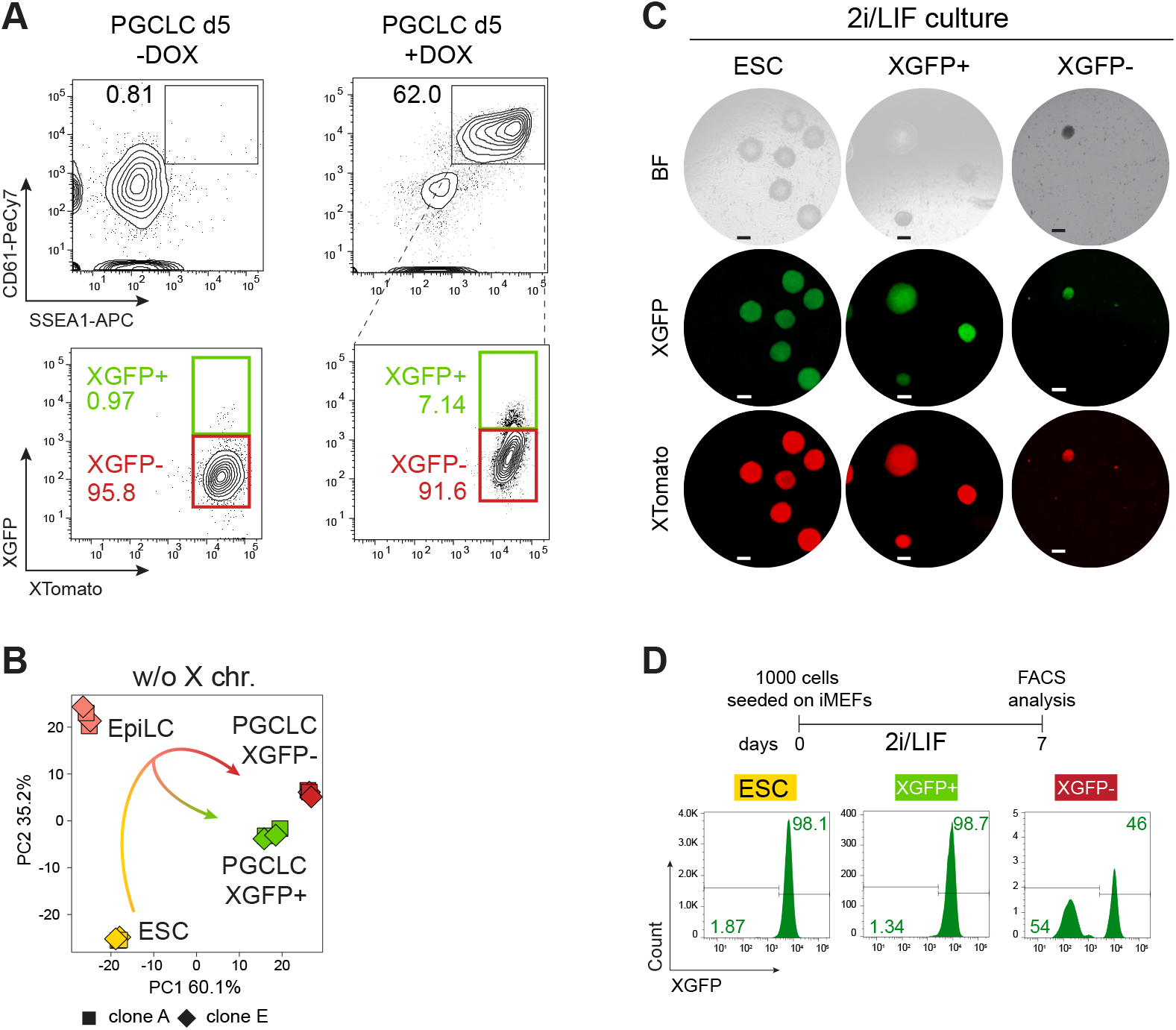
Gene expression analysis reveals two PGCLCs subpopulations. **(A)** Top panel shows representative contour plots of FACS analysis of PGCLC induction, without or with Dox, in PGCLC d5. The number indicates the gated germ cells identified by CD61 and SSEA1 signal. Bottom panel indicates the percentages of X-active (green box) or X-inactive (red box) cells. For the +Dox condition, percentages of X-active and X-inactive originating from SSEA1+/CD61+ cells are shown. Shown are contour plots gated on live cells. **(B)** PCA of gene expression dynamics during PGCLC differentiation. n = top 500 most variable genes excluding X chromosomal genes. Axes indicate the variance. Arrows indicate hypothetical trajectory. Shapes indicate the clones (A11 = square, E9 = rhombus). **(C)** Representative images showing the X-activity reporter status in colonies formed by ESCs, XGFP+ PGCLCs and XGFP-PGCLCs after 7 days of culture in 2i/LIF on immortalized feeder cells. BF = bright field. Scale bar 200 μm. **(D)** FACS analysis showing the X-reporter status of the indicated cell types after 7 days of culture in 2i/LIF on immortalized feeder cells. Numbers indicate the percentage of cells falling in the corresponding gate. Histograms come from XTomato+ gated cells depleted of immortalized feeder cells.

As our transcriptomics and cell cycle analysis suggested that XGFP+ PGCLCs could correspond to an aberrant PGCLC state with similarities to ESCs, we set out to address if this would also lead to an advantage in growth and survival under physiological conditions favoring ground state pluripotent stem cells. We therefore isolated XGFP+ and XGFP-PGCLCs at day 5 and seeded them (1000 cells per 6-well) on irradiated mouse embryonic fibroblasts in 2i/LIF medium (Fig. 2H and I), which previously has been reported to allow the establishment of pluripotent embryonic germ cell (EGC) lines from *in vivo* mouse PGCs (Leitch *et al*, 2010). When we then compared EGC colony formation capacity, we found that while almost no colonies (2 from 1000 seeded cells) originated from XGFP-PGCLCs, we observed a substantially higher number of colonies (84) from XGFP+ PGCLCs, albeit still fewer than when re-plating ESCs (633 colonies). Importantly, both ESCs and XGFP+ PGCLCs retained two active X chromosomes, while only a subset of XGFP-PGCLCs had undergone XGFP-reactivation during EGC colony formation (Fig. EV2C and D).

In summary, RNA expression analysis of XGFP+ and XGFP-PGCLCs showed a PGC-like transcriptome of both populations, further suggesting that X-inactivation and PGCLC induction can be uncoupled in our system. However, we observed that XGFP+ PGCLCs displayed higher expression of several naive pluripotency genes as well as premature expression of a subset of meiotic genes and a rapid cell cycle. Moreover, considering their higher ability to form EGC colonies under ground state pluripotency conditions, this suggests that XGFP+ PGCLCs may correspond to an aberrant PGCLC state of pluripotent stem cell-like character, indicating that X-inactivation could be necessary for correct PGCLC maturation.

### Moderate X-inactivation in XGFP-PGCLCs

To this point, due to the lack of an allele-specific transcriptomic analysis, the X-inactivation and - reactivation dynamics during mouse PGC development *in vivo* and *in vitro* have not been assessed on a chromosome-wide level. Therefore, to determine X chromosome-wide gene inactivation kinetics during PGCLC differentiation, we assessed the allelic expression ratio between the inactive X^mus^ and the active X^cas^. We performed PCA of the allelic ratio of our samples and additionally of neural progenitor cells (NPCs) from the same parental clone (Bauer *et al*, 2021) to include a cell type shown to have undergone complete X-inactivation (Fig. 3A). We observed that the PC1 of the PCA defined the degree of X-inactivation, separating samples with two active Xs on the left (ESCs and XGFP+ PGCLCs), and with one inactive X on the right (XGFP-PGCLCs and NPCs). Moreover, we noticed that EpiLCs were positioned at the center, suggesting an intermediate degree of X-inactivation. We next determined X-inactivation kinetics, while focussing on genes biallelically expressed in ESCs (Fig. EV3A) (allelic expression ratio >0.3 and <0.7) and established an X-inactivation cutoff of an allelic ratio of 0.135, according to the distribution in NPCs (Fig. EV3B). As a control, we assessed the allelic expression ratio of the fully hybrid chromosome 13, which maintained biallelic expression throughout the time course (Fig. EV3C). In contrast, we observed initiation of X-linked gene silencing in EpiLCs, progressing further in XGFP-PGCLCs, while XGFP+ PGCLCs showed biallelic expression, similar to ESCs (Fig. 3B). To assess X-inactivation dynamics in more detail, we grouped X-linked genes according to their silencing kinetics (Fig. 3C). We found 62 genes to have undergone X-inactivation (XCI) in EpiLCs, termed (early XCI), and 138 genes to have undergone inactivation in PGCLCs (late XCI). To our surprise, we observed a large number of genes (93) to still be active in XGFP-PGCLCs (escapees). In comparison, we observed 51 genes escaping X-inactivation in NPCs (Bauer *et al*, 2021), out of which 37 were also found to be escapees in PGCLCs (Fig. 3D). While a certain degree of escape from X-inactivation is expected, the percentage of escapees we observed for PGCLCs here at 32%, is considerably higher than reported for other cell types (Balaton *et al*, 2021; Marks *et al*, 2015; Peeters *et al*, 2014).

**Figure 3.**
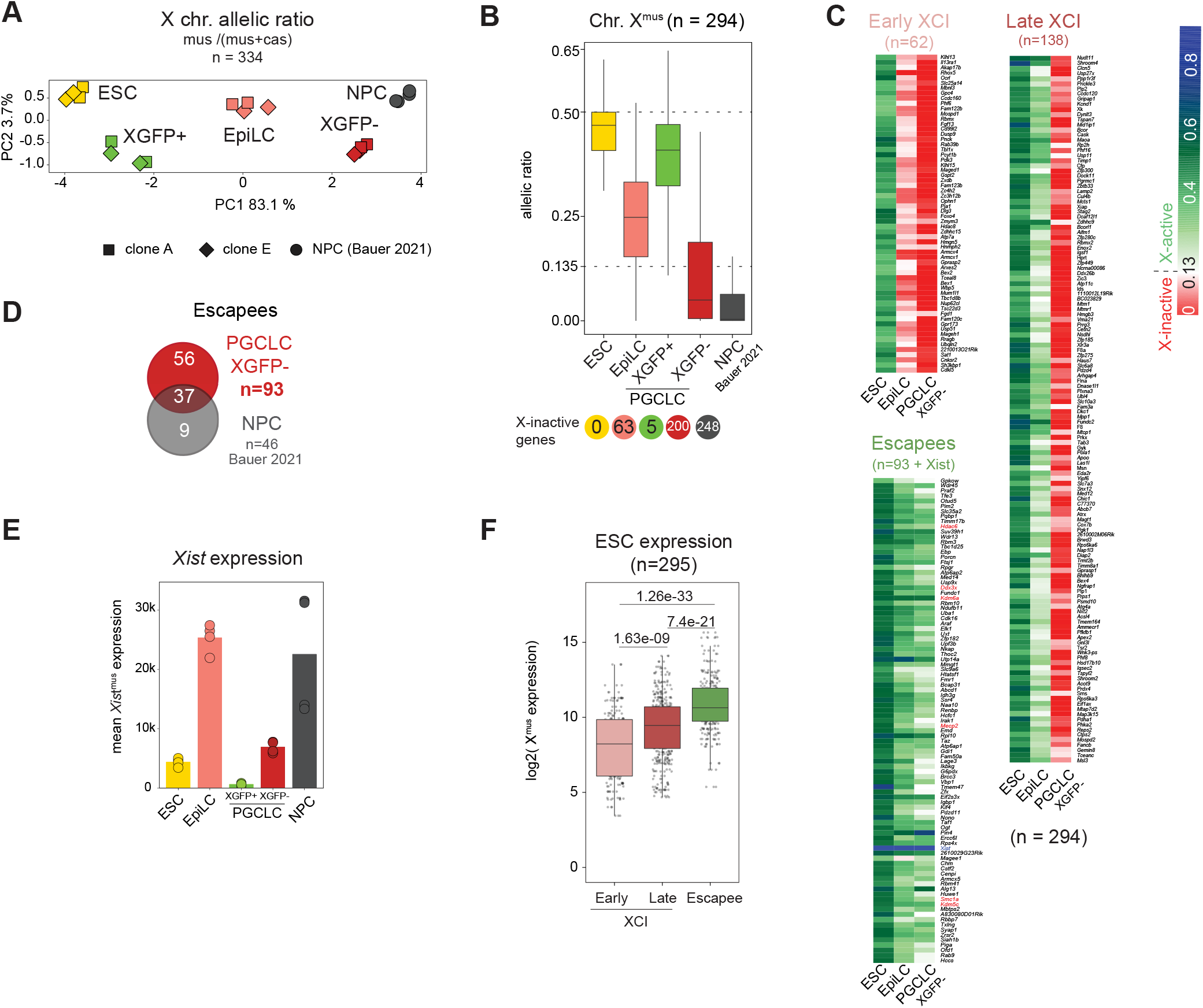
Characterization of X-inactivation dynamics during PGCLC induction. **(A)** PCA of X chromosome allelic ratio (see Methods) for 334 X-linked genes. Axes indicate the variance. Shapes indicate the clones (A11 = square, E9 = rhombus, circle = neural progenitor cells (NPCs) from (Bauer et al, 2021)). **(B)** Boxplots of allelic ratio of X linked genes (n = 294). Upper dashed line indicates biallelic expression with a ratio of 0.5, the lower dashed line indicates the X-inactivation threshold of 0.135. Box plots depict the first and third quartiles as the lower and upper bounds of the box, with a band inside the box showing the median value and whiskers representing 1.5x the interquartile range. Number of X-inactive genes are shown at the bottom. **(C)** Allele-specific expression ratios of X-linked genes are represented as heatmaps, with X-inactive genes in red (ratio ≤ 0.135), X-active genes in green (ratio > 0.135) and mono-allelic Xcas expression in blue (ratio between 0.5 and 1). Genes are ordered by genomic position and subdivided into three groups according to the timing of X-inactivation (early X-inactivation = early XCI, late X-inactivation = late XCI and Escapees). **(D)** Venn diagram showing the total number of Escapee genes overlapping between XGFP-PGCLCs and NPCs from (Bauer et al, 2021). **(E)** Xistmus expression (see Methods). NPCs from (Bauer et al, 2021). Barplot indicates the mean expression value of 4 replicates. **(F)** Expression of Xmus genes in ESCs belonging to the indicated categories. The numbers above the bars indicate p-values (two-sample unpaired Wilcoxon-Mann-Whitney test with R defaults). Box plots depict the first and third quartiles as the lower and upper bounds of the box, with a band inside the box showing the median value and whiskers representing 1.5x the interquartile range. n = 295 genes.

**Figure EV3.**
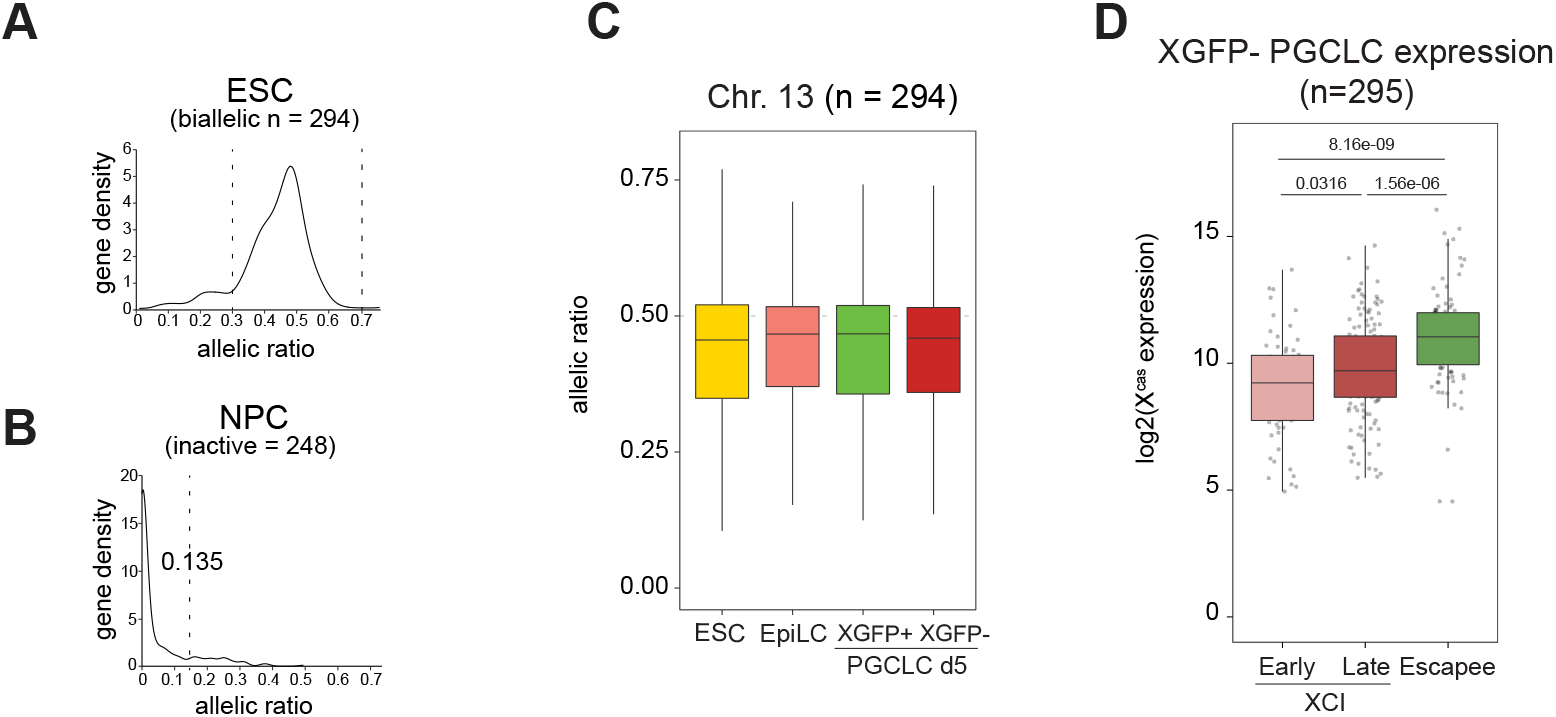
Characterization of X-inactivation dynamics during PGCLC induction. **(A)** Distribution of the allelic ratio of X-linked genes in ESCs. Dashed lines indicate a biallelic expression window from 0.3 to 0.7. n = 334 genes. **(B)** Distribution of the allelic ratio in NPCs of X-linked genes expressed biallelically in ESCs. Dashed line represents allelic ratio of 0.135 used as a threshold for X-inactivation. Genes below the threshold are considered X-inactive. n = 294 genes. **(C)** Boxplots of allelic ratio of genes located on chromosome 13 (n = 294). Dashed line indicates the biallelic ratio of 0.5. Box plots depict the first and third quartiles as the lower and upper bounds of the box, with a band inside the box showing the median value and whiskers representing 1.5x the interquartile range. **(D)** Expression of Xmus genes in XGFP-PGCLCs belonging to the indicated categories. The numbers above the bars indicate p-values (two-sample unpaired Wilcoxon-Mann-Whitney test with R defaults) Box plots depict the first and third quartiles as the lower and upper bounds of the box, with a band inside the box showing the median value and whiskers representing 1.5x the interquartile range. n = 295 genes.

Given these results, we wondered how this large degree of escape from X-inactivation might be explained. We assessed *Xist* expression levels and could observe high levels in EpiLCs, reaching levels comparable to those in NPCs (Fig. 3E). However, expression levels in XGFP-PGCLCs were considerably decreased, which might be explained by the high expression of *Prdm14* in PGCLCs, a known repressor of *Xist* (Payer *et al*, 2013). This is also in line with *in vivo* data (Sugimoto & Abe, 2007), where *Xist* has been shown to be completely downregulated in E10.5 PGCs of equivalent stage (Fig. 2B). Moreover, we wanted to know which features might distinguish escapees from inactivated genes in our system. We measured gene expression levels from the X^mus^ allele in ESCs and found escapees to be significantly higher expressed, while early inactivating genes, in contrast, showed the lowest expression levels (Fig. 3F). Similarly, expression of escapees from the X^cas^ allele was also elevated in XGFP-PGCLCs (Fig. EV3D).

Taken together, we find that PGCLCs undergo a moderate degree of X-inactivation, characterized by a large percentage of escapees. Moreover, our analysis suggests that low expression of *Xist* in PGCLCs might lead to a failure of gene silencing of highly expressed genes, leading to a large percentage of escapees.

### Single-cell RNA-seq analysis of meiotic entry of in vitro-derived germ cells

After having established the degree of X-inactivation during PGCLC specification, we wanted to address the further developmental progression of PGCLCs depending on their X-chromosome status. Having identified and isolated distinct PGCLC types with either two active X-chromosomes (XGFP+ PGCLCs) or one active and one inactivated X-chromosome (XGFP-PGCLCs) (Fig. 2), we were able to assess whether the X-inactivation status of PGCLCs had an impact on germ cell maturation. Furthermore, we sought to investigate to which degree X-reactivation and meiotic entry were intrinsically coupled processes.

To this end, we differentiated XGFP+ and XGFP-PGCLCs using an adapted *in vitro* reconstituted Ovary (rOvary) protocol (Hayashi & Saitou, 2013) and performed single-cell RNA-sequencing (scRNA-seq) using the SMART-Seq v5 Ultra Low Input RNA (SMARTer) Kit for Sequencing (Takara Bio) (Karimi *et al*, 2021). Briefly, we aggregated *in vitro* derived PGCLCs, originating from either XGFP+ or XGFP-populations, for 6 days with somatic cells isolated from E13.5 female embryonic gonads plus mesonephros in order to mimic the female urogenital environment and provide *in vitro*-derived germ cells with the appropriate signaling niche (Hayashi *et al*, 2012; Chuva de Sousa Lopes *et al*, 2008) to facilitate their meiotic entry and X-reactivation (Fig. 4A and B). We then sorted single cells of the following populations on which we performed scRNA-seq. Derived from XGFP-PGCLC rOvaries, we collected three populations: XGFP+ reactivated (144 cells, XTomato+/XGFP+), XGFP intermediate (XTomato+/XGFPint., 144 cells) and XGFP-(XTomato+/XGFP-, 136 cells). From the constitutively active XGFP+ PGCLC rOvaries, we collected one population: XGFP+ constitutive (XTomato+/XGFP+, 188 cells) (Fig. 4A and B). In total, we obtained 391 million reads, with an average of 740,000 reads per cell. Next, to ensure that our analysis focussed on germ cells of appropriate quality, we only included cells with the germ cell marker *Dazl* expression >1 (log2 counts per 10,000) and with sufficient allelic information (see methods). This left us with 379 cells in total and 15,583 informative genes.

**Figure 4.**
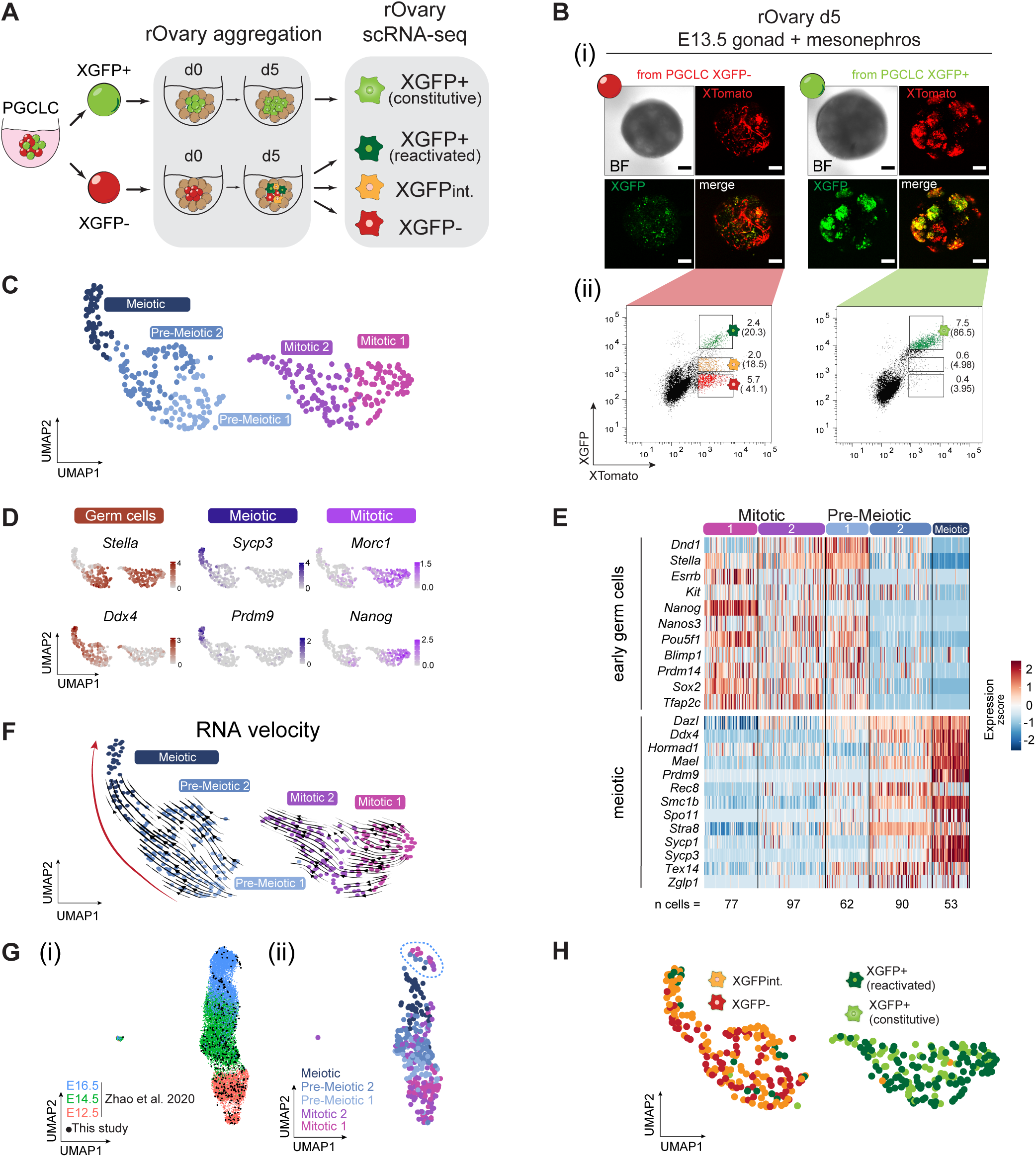
Single-cell RNA-seq of maturing germ cells using the rOvary system. (A) Schematic illustration of the single-cell RNA-seq experiment and the isolated populations during germ cell maturation in rOvaries. The first 24h of culture are indicated as d0. rOvary = reconstituted Ovary, d = day of rOvary culture; XGFP int. = XGFP intermediate. (B) (i) Imaging of XGFP and XTomato reporters in rOvaries d5 aggregated with E13.5 gonadal and mesonephric cells. Scale bars = 50μm. BF = bright field. (ii) FACS gating strategy for single-cells sorted XTomato+ cells against XGFP intensities. Numbers indicate the percentage of gated live cells over the total population. Numbers in brackets indicate the percentage of gated cells over the XTomato+ population. (C) UMAP embedding based on Shared Nearest Neighbor (SNN) modularity clustering identified 5 clusters, termed Mitotic 1 (n = 77), Mitotic 2 (n = 97), Pre-meiotic 1 (n = 62), Pre-meiotic 2 (n = 90) and Meiotic (n = 53) labeled with different colors. (D) Marker gene expression projected onto the UMAP plot. (E) Heatmap of gene expression dynamics throughout germ cell maturation clusters. Selected genes belong to the category “early germ cell” and “meiotic”. Zscore is shown. (F) UMAP of clusters as in (C) with arrows indicating cell trajectories, inferred by RNA velocity analysis. (G) (i) Integration with in vivo published single-cell RNA-seq data from E12.5 (red), E14.5 (green) and E16.5 (blue) (Zhao et al, 2020). Black dots represent cells from in vitro rOvaries from this study. (ii) Distribution of germ cell maturation clusters from rOvaries identified in this study, along the in vivo UMAP projection. Dashed blue circle indicates deviant cells falling in the E16.5 cluster despite not showing expression of late meiotic markers Sycp3 and Prdm9 (data not shown). (H) UMAP projection labeled with FACS sorted populations. XGFPint. = XGFP intermediate.

To characterize cellular heterogeneity, we performed Uniform Manifold Approximation and Projection for dimension reduction (UMAP) on genome-wide single-cell expression data, and then applied Shared Nearest Neighbor (SNN) modularity optimization based clustering which returned 5 clusters (Fig. 4C) that showed distinct patterns according to the expression of mitotic and meiotic germ cell marker genes (Fig. 4D and E). We identified two mitotic clusters termed “Mitotic 1” and “Mitotic 2”, showing expression of the PGC marker *Stella* (also known as *Dppa3*) as well as mitotic PGC markers *Morc1* and *Nanog*. Moreover, we identified two pre-meiotic clusters termed “Pre-meiotic 1” and “Pre-meiotic 2”, defined by the initial expression of both *Stella* and *Ddx4*, and lastly one meiotic cluster termed “Meiotic’’ characterized by expression of the meiotic genes *Prdm9* and *Sycp3*. Next, we wanted to assess whether a directionality within the clusters and eventually among the two groups could be observed. Pseudo-time analysis using RNA velocity (La Manno *et al*, 2018) placed the meiotic cluster at the apex of a path which revealed a differentiation trajectory directed towards meiosis, initiating from the pre-meiotic clusters (Fig. 4F). Moreover, comparison to *in vivo* data (Zhao *et al*, 2020) showed that our mitotic clusters corresponded to E12.5 germ cells, whereas pre-meiotic and meiotic clusters corresponded to later time points; E14.5 and E16.5 (Fig. 4G), confirming that our *in vitro* clusters followed an *in vivo*-like developmental trajectory. Finally, we set out to answer whether our XGFP+ and XGFP-PGCLCs, which were the starting material for our rOvaries (Fig. 4A and B), showed a differential developmental profile, and in particular, if the meiotic germ cells originated preferentially from XGFP+ or XGFP-PGCLCs. To this end, we projected the 4 FACS populations (Fig. 4B and H) on our UMAP plot and observed two major groups, which coincided well with the levels of XGFP fluorescence. One group included predominantly the XGFP-negative and XGFP-intermediate germ cells (originating both from the XGFP-PGCLCs) on the left, and another group was constituted from the XGFP+ reactivated and XGFP+ constitutive germ cells on the right. Intriguingly, both pre-meiotic and meiotic germ cells almost exclusively originated from XGFP-PGCLCs, whereas mitotic germ cells consisted of XGFP+ reactivated and XGFP+ constitutive germ cells.

Taken together, germ cells seem to adopt highly similar transcriptomes when two active X chromosomes are present, irrespective of their parental condition of origin and hence regardless of whether cells underwent X-inactivation followed by X-reactivation, or were constitutively X-active. Moreover, germ cells can undergo X-reactivation in the absence of the meiotic gene expression programme, suggesting that X-reactivation is not dependent on meiotic entry. However, our data suggest that X-inactivation is important for proper germ cell maturation and entry into meiosis, as germ cells originating from constitutively active XGFP+ PGCLCs failed to acquire a meiotic transcriptional profile.

### XGFP-PGCLCs can enter meiotic prophase and undergo oocyte maturation

Our single-cell RNA-seq analysis showed an exclusive ability for XGFP-PGCLCs to differentiate into mature germ cells with a meiotic transcriptional profile. We therefore wanted to further dissect their ability to enter meiosis and their capability to differentiate to more mature stages. To be able to assess in more detail the extent of prophase I progression, we cultured XGFP-PGCLCs for an additional 9 days on immortalized m220 stromal feeder cells in the presence of BMP2 and retinoic acid (Fig. 5A), which was previously shown to facilitate entry into meiosis (Miyauchi *et al*, 2017). During this expansion culture we observed a progressive accumulation of SYCP3+ meiotic cells (Fig. 5B and C), all of which were XGFP+ by day 5 of the expansion culture (Fig. 5B and D), indicating the co-occurrence of XGFP-reactivation with meiotic entry. We then prepared chromosomal spreads from the expansion culture and performed immunostaining for SYCP3, which shows a distinctive pattern according to the different prophase stages (Fig. 5E). Moreover, to aid the correct recognition of the different stages, we stained for the double-strand-break marker, a phosphorylated form of histone variant H2AX (γH2AX) (Mahadevaiah *et al*, 2001). This showed that the majority of cells could successfully enter the zygotene stage by day 9 of expansion culture (Fig. 5E and F), confirming our observation of a meiotic transcriptional profile in cells originating from XFGP-PGCLCs.

**Figure 5.**
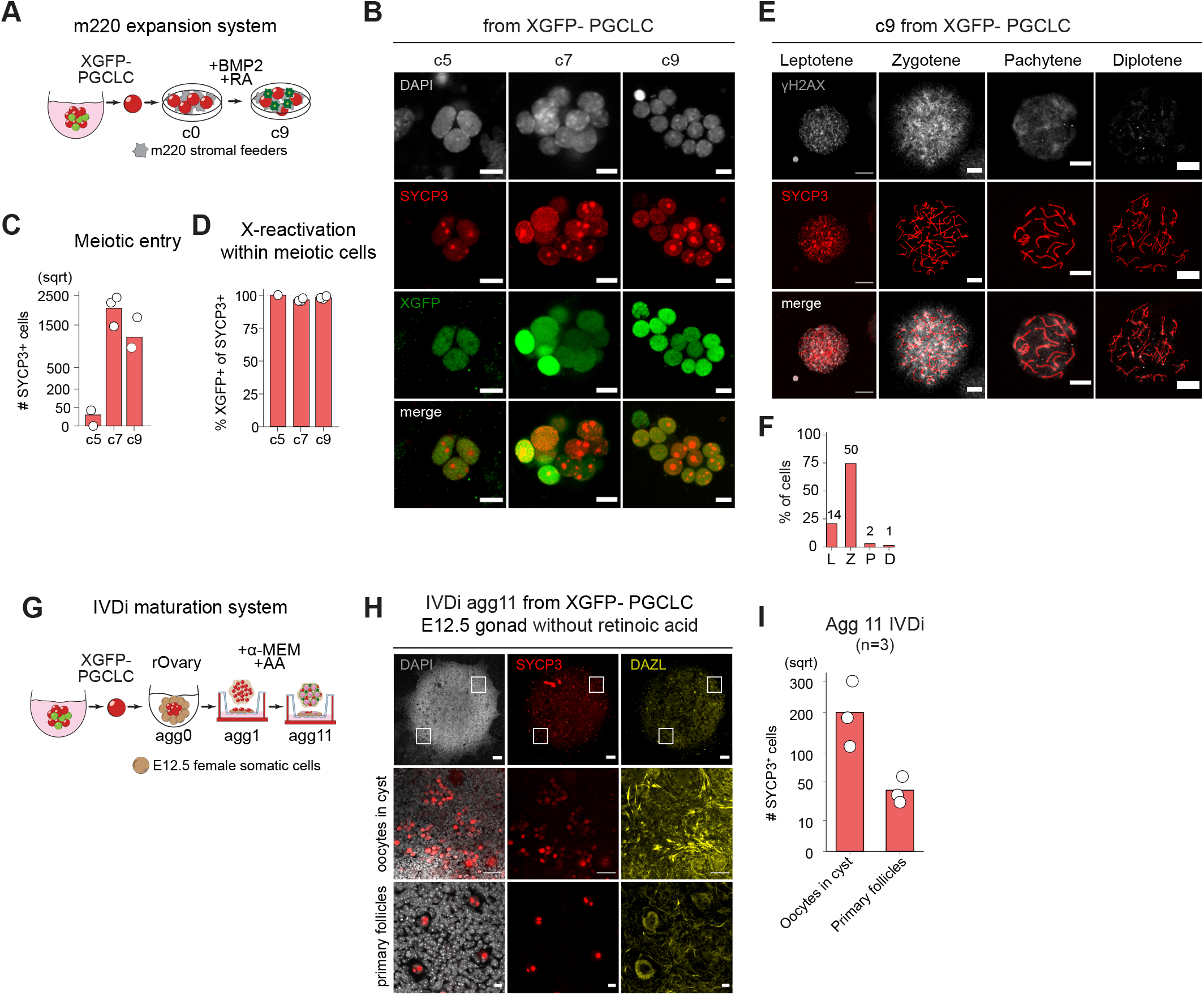
Prophase I progression and germ cell maturation by m220 feeder expansion and IVDi transwell culture. **(A)** Schematic representation of the m220 stromal feeders expansion culture for XGFP-PGCLCs. Meiosis is induced via addition of Retinoic Acid (RA) and Bone Morphogenetic Protein 2 (BMP2). c0 = starting day of culture, c9 = culture day 9 and last day of culture. **(B)** Representative images for the expression of XGFP (green) and SYCP3 (red) in germ cells at c5, c7 and c9 from XGFP-PGCLCs. Cells were counterstained with DAPI (gray). Scale bars = 10 μm. **(C)** Number of SYCP3+ cells per m220 culture day originating from XGFP-PGCLCs. Each white dot represents a biological replicate (n = 3). Y-axis is in square root scale (sqrt) for easier plot visualization. **(D)** Percentage of XGFP+ cells among SYCP3+ cells at the indicated m220 culture day, originating from XGFP-PGCLCs. Each white dot represents a biological replicate (n = 3). **(E)** Representative images showing stages of meiotic prophase I from culture day 9 (c9) germ cells from XGFP-PGCLCs. c9 germ cells were spread and immunostained for SYCP3 (red), and γH2AX (gray). Scale bar = 10 µm. **(F)** Quantification of meiotic progression in culture day (c9) expanded germ cells derived from XGFP-PGCLCs. The graphs show the percentages of the meiotic stage. L, leptotene; Z, zygotene; P, pachytene; D, diplotene. Numbers indicate absolute number of counted cells from n= 1 experiment. **(G)** Schematic representation of the IVDi (In Vitro Differentiation) maturation system. The stages of oogenesis in culture for 11 days are indicated. The condition of culture is indicated above. rOvary = reconstituted Ovary. Agg = aggregation day. AA = Ascorbic Acid. **(H)** Immunofluorescence images of SYCP3 (red), DAZL (yellow) and DAPI at agg11 of IVDi tissue maturation from XGFP-PGCLCs. IVDi = in vitro differentiation. White squares indicate the positions of the magnified section shown below. Top panel scale bar = 100 µm. Middle panel scale bar = 10 µm. Bottom panel scale bar = 50 µm. **(I)** Quantification of SYCP3+ cells (oocytes in cyst and primary follicles) in IVDi tissues at agg11. Each dot represents one IVDi tissue performed in 3 biological replicates.

Next, we wanted to assess if our XGFP-PGCLCs could mature further and initiate oogenesis. We therefore took advantage of a published *in vitro* differentiation protocol and aggregated XGFP-PGCLCs with embryonic-derived somatic gonadal cells, forming a rOvary, followed by the culture of the rOvary onto a transwell to allow *in vitro* differentiation (IVDi) of PGCLCs (Fig. 5G) (Hayashi *et al*, 2017). However, to perform the experiment in a more physiological niche, without external cues, no retinoic acid was added to the IVDi culture and the IVDi tissue was cultured for 11 days until primary follicles had formed. We then stained the entire whole-mount tissue for DAZL and SYCP3 to identify mature (DAZL+) and meiotic (SYCP3+) germ cells and could observe on average 200 oocytes in cysts per aggregate and moreover around 50 primary follicles (Fig. 5H and I), similar to what has been observed previously (Hamada *et al*, 2020).

Taken together, our XGFP-PGCLCs are able to enter prophase I and to mature into primary follicles when defined culture conditions are provided.

### X-Reactivation occurs progressively during germ cell maturation and meiotic entry

Having shown that cells could undergo X-reactivation in the absence of meiosis, we nevertheless wanted to assess if X-reactivation was a prerequisite for meiotic entry. To therefore analyse X-reactivation dynamics in more detail, we again took advantage of the hybrid background of our XRep cell line and performed allele-specific RNA expression analysis, which allowed us to successfully detect allele-specific expression of 220 X-linked genes (see methods). To first assess the X-status on a chromosome-wide level, we calculated the average allelic ratio of all X-linked genes (Fig. 6A). As expected, we observed biallelic X-linked gene expression of the XGFP+ mitotic clusters 1 and 2, as reflected by an average allelic ratio of 0.5. However, cells of the pre-meiotic and meiotic clusters, despite originating from mostly XGFP-and XGFPint. populations showed close to biallelic expression at an average allelic ratio of ∼0.4, as the sensitivity of the XGFP reporter was insufficient to mark cells as reactivated if they had low levels of X-inactivation (Fig. 6B). We therefore assessed the X-status on a gene by gene level and compared it to the data of ESCs, EpiLCs and XGFP-PGCLCs (Fig. 6C and D). In addition to 78 escapees, being active throughout the differentiation, we observed early X-chromosome reactivation (early XCR) of 58 genes in pre-meiotic cells. Therefore, the vast majority of genes (85%) had escaped X-inactivation in the first place, or undergone reactivation, before the onset of meiosis. Moreover, 17 genes reactivated as cells underwent meiosis (late XCR), with only 8 genes still being inactive in meiotic cells (no XCR). Furthermore, we observed that early reactivating genes displayed higher allelic ratios in XGFP-PGCLCs compared to late reactivating genes (Fig. 6D and E), suggesting that the degree of silencing could influence X-reactivation timing.

**Figure 6.**
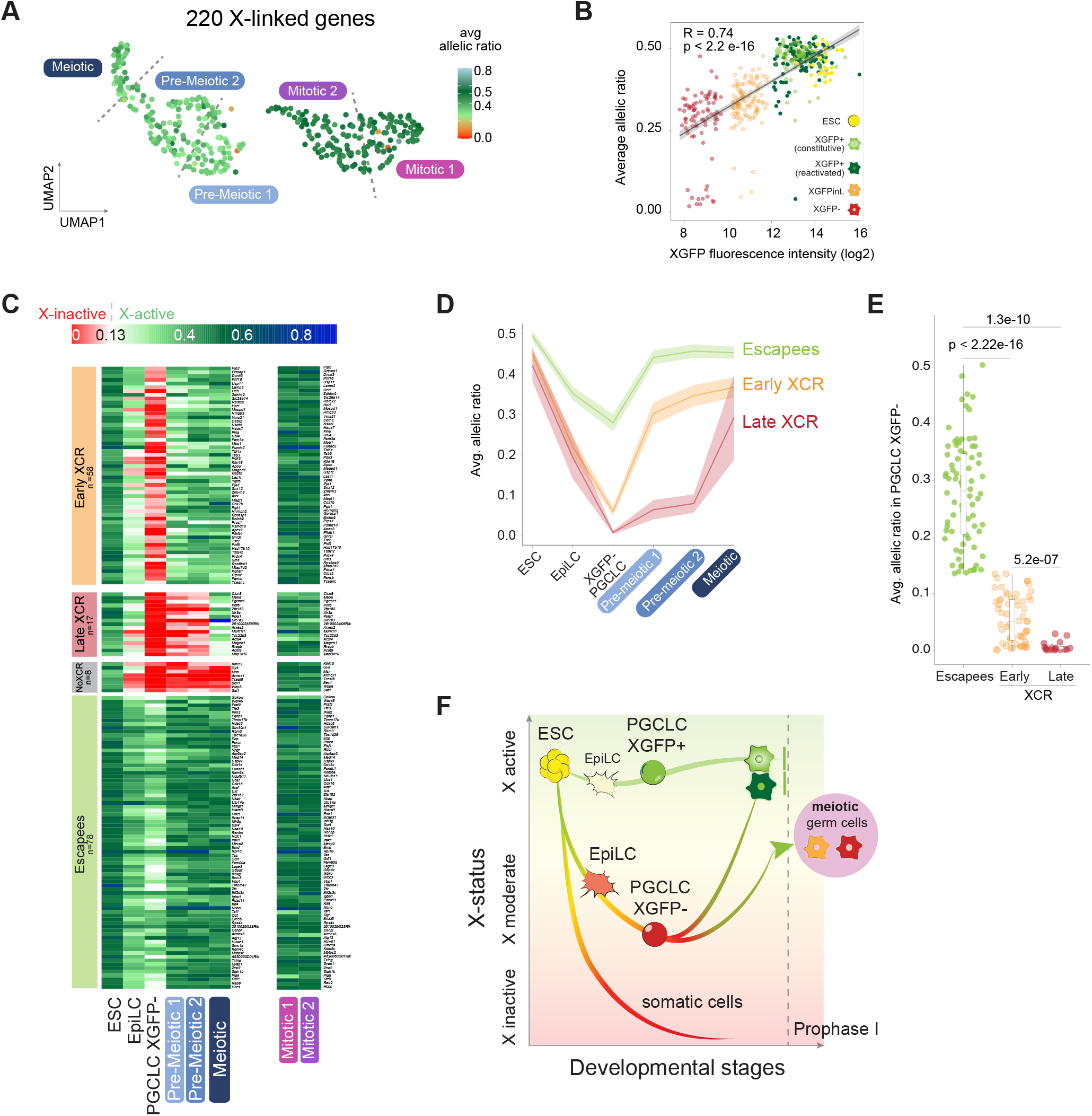
Biallelic re-expression of X-linked genes. **(A)** Average allelic ratio of single cells projected onto the UMAP plot. n = 220 X-linked genes per single cell. X-inactivation (average ratio <0.135) in red and X-reactivation (average ratio from 0.135 > 0.8) in green. Labels indicate the 5 different previously identified clusters. Dashed lines indicate the position of cluster borders. **(B)** Distribution of single cells based on fluorescence intensity of XGFP reporter quantified by BD FACSDiva Software, plotted against the X chromosome average allelic ratio per cell. R and p-values calculated by Pearson’s correlation are shown. Black line represents linear regression fitting. **(C)** Heatmaps of allele-specific ratios of X-linked genes in ESC, EpiLC, XGFP-PGCLC, Pre-meiotic, Meiotic and Mitotic clusters. X-inactive genes are shown in red (ratio ≤ 0.135), X-active genes in green (ratio > 0.135) and mono-allelic Xcas expression in blue (ratio between 0.5 and 1). Colour gradients used in between and above these two values as shown in the legend. Genes are ordered by genomic position and grouped according to the category to which they belong, indicated on the left side of the heatmap. n = 161 genes. **(D)** Average allelic ratios of X-linked genes within each category (Escapees, Early XCR, Late XCR) in ESC, EpiLC, XGFP-PGCLC, Pre-meiotic and Meiotic clusters. **(E)** Each dot indicates the average allelic ratio of a single X-linked gene belonging to the indicated category in XGFP-PGCLCs. The numbers above the bars indicate p-values (two-sample unpaired Wilcoxon-Mann-Whitney test with R defaults). Box plots depict the first and third quartiles as the lower and upper bounds of the box, with a band inside the box showing the median value and whiskers representing 1.5x the interquartile range. **(F)** Working model of the effects of X-status on germ cell developmental stages.

Taken together, X-reactivation in germ cells seems to occur in two waves. First, before the onset of meiosis for the majority of genes and second, concomitantly with meiotic entry for a small subset of genes.

## Discussion

While X-chromosome inactivation has been a long-studied phenomenon (Lyon, 1961) and has been shown to play an important biological role for embryonic development (Marahrens *et al*, 1997) and pluripotency exit (Schulz *et al*, 2014), its reversal by X-reactivation and its biological function during germ cell development have remained elusive to date. Previous studies on X-chromosome dynamics during female mouse germ cell development have been hampered by a lack of allelic-resolution, a low number of genes assessed, as well as an inability to directly trace the X-chromosome status of single cells (Sugimoto & Abe, 2007; Chuva de Sousa Lopes *et al*, 2008). To overcome these limitations, we generated with XRep an *in vitro* system, which allowed us to reveal the X-chromosome inactivation and reactivation cycle and its functional relation to germ cell development and meiotic progression. We thereby uncovered that X-inactivation is an important hallmark of proper PGCLC differentiation in order to progress at later stages towards meiotic entry (Fig. 6F). X-reactivation, on the other hand, coincides temporally with meiotic maturation. This is in line with the timing of X-reactivation in mouse germ cells *in vivo*, (Sugimoto & Abe, 2007; Sangrithi *et al*, 2017; Chuva de Sousa Lopes *et al*, 2008), where it takes place gradually, initiating during germ cell migration and peaking after colonization of the gonads around the time of meiotic entry. Additionally, our *in vitro* system enabled the isolation of PGCLCs harboring two active X, a unique advantage over *in vivo* systems, as it allowed us to compare the differentiation potential of PGCLCs with and without X-inactivation. While our results suggest that PGCLC specification can occur in the absence of X-inactivation, we found that germ cells, which had never undergone X-inactivation in the first place, or in which X-reactivation occurred preemptively, displayed a mitotic germ cell character and did not enter a meiotic trajectory. This further highlights how timely X-inactivation and -reactivation might be necessary for proper germ cell maturation (Fig. 6F). Moreover, while we acknowledge that our findings are based on data generated *in vitro*, we note that allele-specific single-cell RNA-seq of E5.5-E6.5 epiblast cells, the precursors of PGCs, revealed a considerable heterogeneity in X-inactivation progression at this developmental time window (Mohammed *et al*, 2017; Lentini *et al*, 2021; Naik *et al*, 2021; Cheng *et al*, 2019), which could potentially allow cells to give rise to XaXa PGCs, similar to our XGFP+ PGCLCs. Thus our data supports the idea of a potential functional link between appropriate X-chromosome dosage compensation kinetics and developmental progression during mammalian germ cell maturation.

It remains an open question, what could be the potential role of X-inactivation for proper PGCLC development and if it is a driver or, alternatively, a diagnostic mark for meiotic competence of germ cells. We observed that XGFP+ PGCLCs, which failed to undergo X-inactivation, differed from XGFP-PGCLCs on multiple accounts. Albeit sharing an overall similar transcriptome signature with their XGFP-germ cell counterparts, XGFP+ PGCLCs displayed ESC-like features including a higher expression of naive pluripotency genes, shortened cell cycle and propensity to form pluripotent EGC colonies when cultured under 2i/LIF conditions. An explanation for this pluripotency-related phenotype could be the two-fold expression of critical X-linked dosage-sensitive genes, which need to be silenced by X-inactivation to allow normal pluripotency exit during ESC differentiation (Schulz *et al*, 2014). For example, *Dusp9*, an X-linked regulator of MAPK signaling, has been shown to be responsible for the lower DNA-methylation levels of XX pluripotent stem cells, when compared with XY and XO cells (Choi *et al*, 2017; Song *et al*, 2019; Genolet *et al*, 2021). In germ cell development, DNA methylation safeguards repression of late germ cell / meiotic genes during early germ cell stages and demethylation of their promoters is required for their upregulation during germ cell maturation and meiotic entry (Hill *et al*, 2018; Yamaguchi *et al*, 2012). Along those lines, we observed that XGFP+ PGCLCs also displayed precocious expression of a subset of late germ cell markers, which remained repressed in XGFP-PGCLCs. Importantly, demethylation of late germ cell genes alone has been shown to only lead to partial activation of some germ cell genes, while not being sufficient for their full expression in the absence of meiosis-inducing signals (Ohta *et al*, 2017; Miyauchi *et al*, 2017). This would explain our observation of a relatively mild upregulation of late germ cell genes in our XGFP+ PGCLCs and why this was not sufficient to aid entrance of XGFP+ cells into a full meiotic trajectory after their aggregation with gonadal somatic cells.

*Klhl13*, another X-linked MAPK pathway regulator, has been recently described to promote pluripotency factor expression, thereby delaying differentiation when expressed at double dose (Genolet *et al*, 2021). The counterbalance between pluripotency vs. differentiation-promoting signaling responses was also observed in our gene expression analysis, in which we found “MAPK regulation” and “WNT signaling” to be enriched GO terms in XGFP-PGCLCs, while “response to LIF” was enriched in XGFP+ PGCLCs (Fig. 2F). Apart from being involved in pluripotency, MAPK-inhibition (Kimura *et al*, 2014) as well as WNT- and LIF-signaling pathways (Ohinata *et al*, 2009; Hayashi *et al*, 2011) play facilitating roles during PGCLC induction, therefore differential enrichment of these pathways in our XGFP+ and XGFP-PGCLCs might contribute to their distinct developmental potentials. Taken together, the combination of these differential features might lead to a reduced mitotic propensity of XGFP-PGCLCs, which might prime them for meiotic entry, while XGFP+ PGCLCs rather remain mitotic and do not enter meiosis. To which degree this may be a cause or consequence of the X-inactivation status in PGCLCs and how X-linked gene dosage might affect female germ cell development will need to be addressed by future studies.

While we found that X-inactivation marked PGCLCs of full potential for subsequent meiosis and oogenesis, X-reactivation occurred progressively during their transition from pre-meiotic into meiotic stages. Evidently, X-reactivation is not dependent on meiotic entry as it occurred completely in mitotic germ cells as well, and X-reactivation by itself was also not sufficient for germ cells to enter a meiotic trajectory. However, it remains to be tested whether X-reactivation is a requirement for female germ cells to progress through meiosis, or if the two processes are functionally unrelated. As in the case of pluripotency, reactivation of dosage-sensitive X-linked genes could enable the initiation of the meiotic gene expression program by promoting the derepression and upregulation of meiotic genes (Hill *et al*, 2018; Yamaguchi *et al*, 2012). The absence of double X dosage and/or abnormalities in meiotic pairing ability greatly diminishes the success rate of XO and XY germ cells to pass through meiotic prophase due to delay of meiotic initiation and meiotic arrest when compared to XX germ cells (Hamada *et al*, 2020). Therefore, equalizing the chromatin state between the heterochromatic inactive X and euchromatic active X by X-reactivation could be a necessary step in order to allow X-X chromosome pairing during meiotic prophase. Our XRep system will provide a unique tool to test the potential requirement of X-reactivation for meiotic progression and thereby reveal the biological function of the intriguing epigenetic yoyo of X-inactivation and -reactivation in the mammalian germ cell lineage.

## Materials and Methods

### Cell culture

#### Embryonic stem cell culture: Serum/LIF

Embryonic Stem Cells (ESCs) were maintained and expanded on 0.2% gelatin-coated dishes in DMEM (Thermo Fisher Scientific, 31966021) supplemented with 10% Fetal Bovine Serum (FBS) (ES-qualified, Thermo Fisher Scientific, 16141079), 1,000 U/ml LIF (ORF Genetics, 01-A1140-0100), 1 mM Sodium Pyruvate (Thermo Fisher Scientific, 11360070), 1x MEM Non-Essential Amino Acids Solution (Thermo Fisher Scientific, 11140050), 50 U/ml penicillin/streptomycin (Ibian Tech, P06-07100) and 0.1 mM 2-mercaptoethanol (Thermo Fisher Scientific, 31350010). Cells were cultured at 37°C with 5% CO_2_. Medium was changed every day and cells were passaged using 0.05% Trypsin-EDTA (Thermo Fisher Scientific, 25300054) and quenched 1:5 in DMEM supplemented with 10% FBS (Life Technologies, 10270106). Cells were monthly tested for mycoplasma contamination by PCR.

#### Embryonic stem cell culture: 2i/LIF

ESCs were cultured for 24h prior to the start of the primordial germ cell-like cell induction in 2i/LIF medium. Briefly, a homemade version of the N2B27 medium was prepared based on previous reports (Ying *et al*, 2008) with additional modifications reported in (Hayashi & Saitou, 2013) containing two chemical inhibitors 0.4 µM PD032591 (Selleck Chemicals, S1036) and 3 µM CHIR99021 (SML1046, SML1046) together with 1,000 U/ml LIF (ORF Genetics, 01-A1140-0100). ESCs were seeded on a dish coated with 0.01% poly-L-ornithine (Sigma-Aldrich, P3655) and 500 ng/ml laminin (Corning, 354232).

#### XRep cell line generation

We used the female F2 ESC line EL16.7 TST, derived from a cross of *Mus musculus musculus* with *Mus musculus castaneus* (Ogawa *et al*, 2008). As a result, cells contain one X chromosome from *M*.*m musculus* (X^mus^) and one from *M*.*m castaneus* (X^cas^). Moreover, EL16.7 TST contains a truncation of Tsix on X^mus^ (Tsix^TST/+^), which abrogates *Tsix* expression and leads to the non-random inactivation of X^mus^ upon differentiation. XGFP and XtdTomato vectors were integrated first, followed by integration of rtTA and last of germ cell transcription factor vectors.

#### XGFP and XtdTomato dual color reporter

A GFP reporter construct (Wu *et al*, 2014) was targeted in the second exon of *Hprt* on X^mus^ as described in (Bauer *et al*, 2021). The same strategy was used to simultaneously target a tdTomato reporter construct in the second exon of *Hprt* on X^cas^ and a GFP reporter on X^mus^. Briefly, 5×10^6^ EL16.7 TST ESCs were nucleofected with the AMAXA Mouse Embryonic Stem Cell Nucleofector Kit (LONZA, VPH-1001) using program A-30 with 1.6 µg each of GFP and tdTomato circularised targeting vectors and 5 µg single gRNA vector PX459 (5’-TATACCTAATCATTATGCCG-3’) (Addgene, 48139, a gift from Feng Zhang). Homology arms flanking the target site were amplified from genomic DNA and cloned into pBluescript II SK(+) (Addgene, 212205) by restriction-enzyme based cloning and the cHS4-CAG-nlstdTomato-cHS4 and cHS4-CAG-nlsGFP-cHS4constructs, kindly provided by J. Nathans (Wu *et al*, 2014) were cloned between the two homology arms. 7.5 µM of RS-1 (Merck, 553510) was added to enhance homology-directed repair. To select for the homozygous disruption of *Hprt*, cells were grown in the presence of 10 µM 6-thioguanine (Sigma-Aldrich, A4882-250MG) for 6 days, and GFP+ / tdTomato+ cells were isolated by FACS using a BD Influx (BD Biosciences). Single clones were screened by Southern blot hybridization as described in (Bauer *et al*, 2021).

#### Rosa26 rtTA

1 µg of R26P-M2rtTA targeting vector (Addgene, 47381) and 5 µg of PX459 gRNA vector (5’-GACTCCAGTCTTTCTAGAAGA-3’) were nucleofected with the AMAXA Mouse Embryonic Stem Cell Nucleofector Kit (LONZA, VPH-100) using program A-30 in the XRep. Cells were selected with 3 μg/ml puromycin (Ibian tech., ant-pr-1) for 5 days, with medium being changed daily. Single clones were screened for rtTA expression by quantitative RT-PCR and by Southern blot hybridization, with genomic DNA being digested by EcoRV.

#### Germ cell transcription factors overexpression

PB-TET vectors containing key germ cell factors Blimp1, Tfap2c and Prdm14 (Nakaki *et al*, 2013) were kindly given by F. Nakaki. Cells were transfected with 3 µg each of PB-TET vectors, pPBCAG-hph and a PiggyBac Transposase vector using the AMAXA Mouse Embryonic Stem Cell Nucleofector Kit (LONZA, VPH-1001). Transfected cells were selected with 200 μg/ml hygromycin B Gold (Ibian tech., ant-hg-1) for 10 days and genotyped by PCR for transgenes. The primer sequences are shown in Table 1.

**Table 1.**
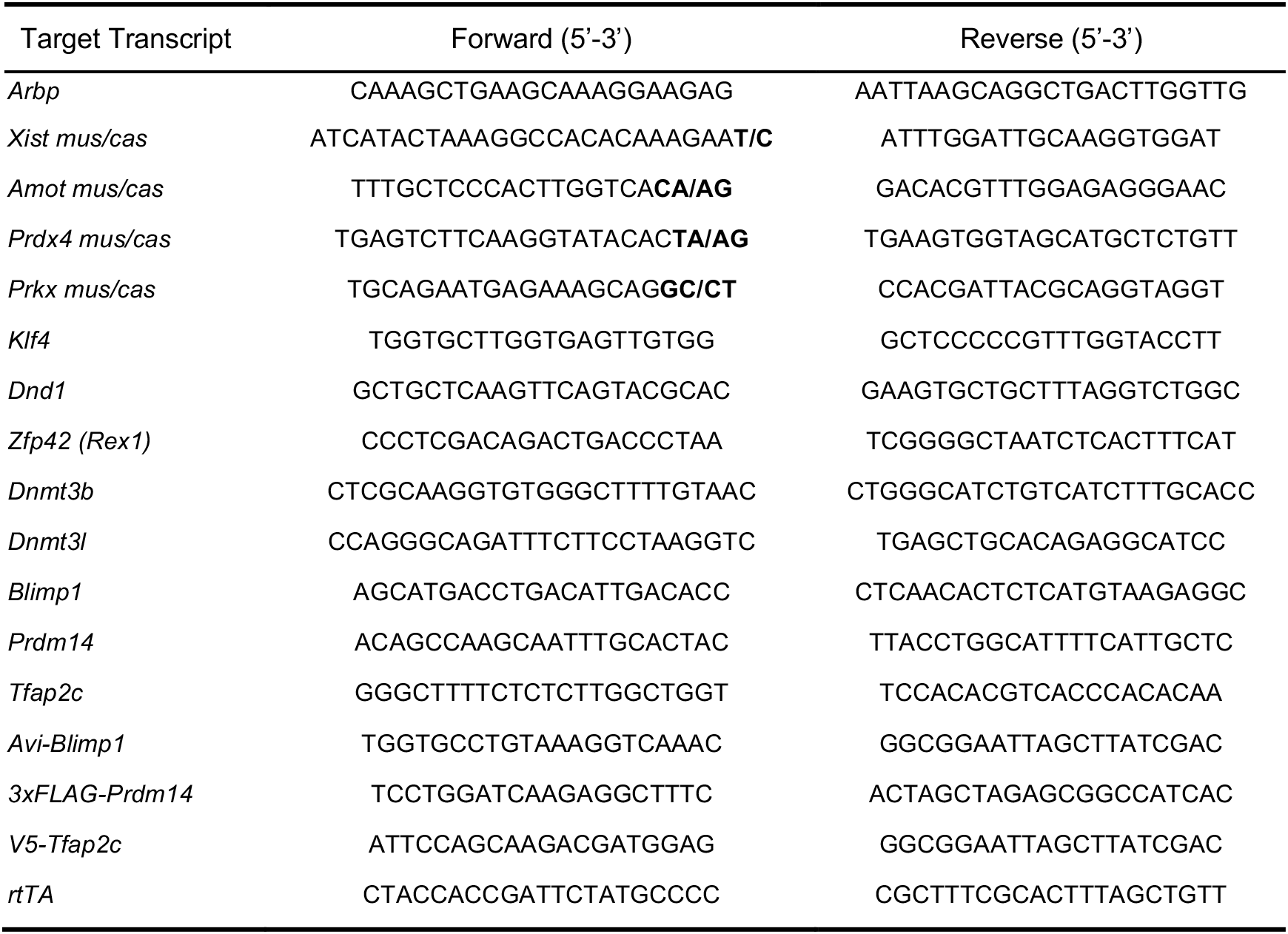
Primer sequences used in this study.

Copy number integration was estimated by Southern blot hybridization. Briefly, 15 µg of genomic DNA were digested with BamHI. DNA fragments were electrophoresed in 0.8% agarose gel and transferred to an Amersham Hybond XL membrane (GE Healthcare, RPN303S). The b-geo probe was designed downstream of the BamHI site, obtained by digesting the PB-TET-Avi-Blimp1 plasmid with CpoI/SmaI, labeled with dCTP [α-32P] (Perkin Elmer, NEG513H250UC) using High Prime (Roche, 11585592001), purified with an illustra ProbeQuant G-50 Micro Column (GE Healthcare, 28903408) and hybridization performed in Church buffer. Radioisotope images were captured with a Phosphorimager Typhoon Trio

### Epiblast-like cell and primordial germ cell-like cell induction

XRep ESCs were induced into PGCLCs as described previously (Hayashi & Saitou, 2013) with the following modifications as this condition was most efficient in generating PGCLCs. ESCs were thawed on 0.2% gelatin in serum/LIF and after 24h seeded at a density of 0.6 x10^5^ cells/cm^2^ in 2i/LIF medium on a dish coated with 0.01% poly-L-ornithine (Sigma-Aldrich, P3655) and 500 ng/ml laminin (Corning, 354232). 24h later, ESCs were dissociated with TrypLE Express for 5 mins at 37°C and induced into EpiLCs by addition of human recombinant basic fibroblast growth factor (bFGF) (Invitrogen, 13256-029) and activin A (Peprotech, 120-14P) and seeding on 16.7 µg/ml human plasma fibronectin-coated plates (Merck Millipore, FC010). After 48h, EpiLCs were split using TrypLE Express (Life Technologies 12604013) and re-seeded at 0.2 x 10^5^ cells/cm^2^ on 16.7 µg/ml human plasma fibronectin-coated plates. After an additional 48h, EpiLCs were aggregated in U-bottom 96-well Lipidure-Coat plate (Thermo Fisher Scientific, 81100525) at 2,000 cells per aggregate in GK15 medium (GMEM (Life Technologies, 11710035), 15% KnockOut Serum Replacement (KSR) (Thermo Fisher, 10828028), 0.1 mM nonessential amino acids (NEAA) (Thermo Fisher Scientific, 11140050), 1 mM sodium pyruvate (Thermo Fisher Scientific, 11360), 2 mM Glutamax (Life Technologies, 35050061), 0.1 mM 2-mercaptoethanol (Thermo Fisher Scientific, 21985-023), and 100 U/ml penicillin and 0.1 mg/ml streptomycin (Thermo Fisher Scientific, 15140) with 1.5 µg/ml doxycycline (Tocris, 4090/50) for 5 days.

#### PGCLCs mitotic expansion

PGCLC mitotic expansion culture was performed as previously described (Ohta *et al*, 2017) with few modifications. Briefly, five days after PGCLC induction, SSEA1+/CD61+ PGCLCs were sorted by flow cytometry onto m220 feeder cells, which constitutively express a membrane-bound form of mouse Stem Cell Factor (Dolci *et al*, 1991; Majumdar *et al*, 1994) on 0.1% gelatin-coated optical bottom plates (Nunc, 165305). The expansion culture was maintained for a total of 9 days. The first 3 days in GMEM containing 100 ng/ml SCF (Peprotech, 250-03), 10 µM forskolin (Sigma-Aldrich, F3917), 10 µM rolipram (Abcam, ab120029), 2.5% FBS (Capricorn Scientific, FBSES12B), 10% KSR, 0.1 mM NEAA, 1 mM sodium pyruvate, 2 mM Glutamax (Life Technologies, 35050061), 0.1 mM 2-mercaptoethanol, 100 U/ml penicillin, 0.1 mg/ml streptomycin and 100 nM all-trans Retinoic Acid (RA) (Enzo Life Sciences, BMLGR100).

#### PGCLCs meiosis induction

Meiosis was induced after 3 days of mitotic expansion culture as previously reported (Miyauchi *et al*, 2017, 2018) by a combined treatment of 300 ng/ml BMP2 (R&D Systems, 355-BM) and 100 nM RA. Medium was replaced completely every two days until the end of the culture period.

#### rOvary reconstitution

10,000 sorted SSEA1+/CD61+ PGCLCs were mixed with 75,000 freshly thawed E13.5 female somatic gonadal and mesonephric cells (SSEA1-/CD31-) or E12.5 female somatic gonadal cells and cultured in Lipidure-Coat plates at 37°C in a 5% CO_2_ incubator for 6 days for the scRNAseq protocol or for 2 days for the IVDi as described in (Hayashi *et al*, 2017). Mouse care and procedures were conducted according to the protocols approved by the Ethics Committee on Animal Research of the Parc de Recerca Biomèdica de Barcelona (PRBB) and by the Departament de Territori i Sostenibilitat of the Generalitat de Catalunya.

#### Oocyte *in vitro* differentiation (IVDi) culture

IVDi culture was performed as previously described (Hayashi *et al*, 2017). Briefly, one single rOvary was placed in the middle of a 24-well Transwell-COL membrane (Corning, CLS3470-48EA) and cultured in alpha-MEM (Life Technologies, 12571063) with 0.15 mM ascorbic acid (Sigma-Aldrich, A7506), 2% FBS, 2 mM Glutamax (Life Technologies, 35050061), 0.1 mM 2-mercaptoethanol, 50 U/ml penicillin/streptomycin under normoxic condition (20% O_2_ and 5% CO_2_ at 37°C) for 11 days, changing IVDi medium every other day.

#### Fluorescence-activated cell sorting (FACS)

After 5 days of culture, PGCLC aggregates were dissociated using TrypLE Express (Thermo Fisher Scientific, 12604021) for 8 min at 37°C, with periodical tap-mixing. The reaction was quenched 1:5 with wash buffer DMEM/F12 (Thermo Fisher Scientific, 11320-082) containing 0.1% bovine serum albumin (BSA) fraction V (Thermo Fisher Scientific, 15260-037) and 30 mM HEPES (Gibco, 15630-056) containing 0.1 mg/mL of DNAse I (Sigma-Aldrich, DN25-10MG). The cell suspension was centrifuged at 1200 rpm for 5 min, resuspended in FACS buffer (0.1% BSA in PBS) and passed through a 70 µm cell strainer (Corning, 352350). Cells were stained with 1:100 SSEA1-eFluor 660 (Thermo Fisher Scientific, 50-8813-42) and 1:10 CD61-PE-Vio770 (Miltenyi Biotec, 130102627) for 1h at 4°C. Cells were washed thrice with FACS Buffer, stained with 1:1000 DAPI (Thermo Fisher Scientific, D1306) and then FACS sorted using a BD FACSAria II or a BD Influx. Double-positive population of PGCLCs was collected in GK15 medium. Data was analysed with Flowjo (Tree Star) software.

#### Cell cycle analysis

Identification of G1, S, G2/M cell cycle phases was based on DNA content and performed as described previously (Bonev *et al*, 2017) with minor modifications. Briefly, ESCs, EpiLC, PGCLCs were dissociated and quenched as described above. Cells were then fixed for 10 min at room temperature with freshly prepared 1% formaldehyde in PBS (Sigma-Aldrich, F8775-4×25ML) and the reaction then quenched by addition of 0.2M glycine (NZYTech, MB01401) 15 min on ice. 1×10^6^ cells/ml were permeabilized using 0.1% saponin (Sigma-Aldrich, 47036-50G-F) containing 10 µg/ml DAPI (Thermo Fisher Scientific, D1306) and 100 µg/ml RNase A (Thermo Fisher Scientific, EN0531) for 30 min at room temperature protected from light with slight agitation. After washing once with cold PBS, samples were resuspended in cold 0.5% BSA in PBS at a concentration of 1×10^6^ cells/ml and immediately analyzed using a BD LSRFortessa.

#### Immunofluorescence of PGCLCs bodies and rOvaries

Immunofluorescence analysis of PGCLC bodies or rOvaries was performed on cryosections prepared as follows: aggregates were fixed with 4% paraformaldehyde (PFA) (Electron Microscopy Science, 15713) in PBS at room temperature for 30 min, followed by three washes in PBS and submerged in serial concentrations of 10% and 30% of sucrose (Sigma-Aldrich, S0389) in PBS, 15 mins and overnight at 4°C respectively. The samples were embedded in OCT compound (Sakura Finetek, 4583), snap-frozen in liquid nitrogen, and cryo-sectioned at a thickness of 10 µm at −20°C on a cryostat (Leica, CM1850). The sections were placed on a coated glass slide (MAS-GP type A; Matsunami, S9901) and dried completely.

For immunostaining, the slides were blocked with PBS containing 10% normal goat serum (NGS) (Abcam, ab7481), 3% BSA (Sigma-Aldrich, A3311), and 0.2% Triton X-100 (Sigma-Aldrich, T9284) for 1 hr at room temperature, followed by incubation with the primary antibodies diluted in a 1:1 solution of blocking buffer to PBS with 0.2% Tween (PBST) (Sigma-Aldrich, P7949) overnight at room temperature. The slides were washed three times with PBST, then incubated with the secondary antibodies diluted as the primary, with DAPI at 1 µg/ml for 1 hr at room temperature. Following three washes in PBST, the samples were mounted in VECTASHIELD with DAPI (Vector Laboratories, H1200) and observed under a Leica SP8 confocal microscope. All images were analyzed using Fiji/Image J software (Schindelin *et al*, 2012). All antibodies used in this study are listed in Table 2.

**Table 2.** Antibodies used in this study.

| Name | Description | Dilution | Company | Catalog# |
| --- | --- | --- | --- | --- |
| **Primary antibody** |  |  |  |  |
| anti-Sox2 | Rabbit polyclonal | 100x | Abcam | ab97959 |
| anti-Tfap2 (6E4/4) | Mouse monoclonal | 300x | Santa Cruz | SC12762 |
| anti-cleaved PARP1 | Rabbit monoclonal | 100x | Abcam | ab32064 |
| anti-Sycp3 | Mouse monoclonal | 100x | Abcam | ab97672 |
| anti-γH2A.X S139 | Rabbit polyclonal | 100x | Abcam | ab11174 |
| anti-Dazl | Rabbit polyclonal | 200x | Abcam | ab34139 |
| anti-GFP | Chicken polyclonal | 500x | Abcam | ab13970 |
| **Surface markers** |  |  |  |  |
| SSEA1-eFluor 660 | Mouse monoclonal | 50x | Thermos | 50-8813-42 |
| CD61-PE-Vio770 | Hamster monoclonal | 10x | Miltenyi Biotec | 130-102-627 |
| **Secondary antibody** |  |  |  |  |
| Anti-chicken IgY | Goat polyclonal / Alexa488 | 500x | Life Technologies | A11039 |
| Anti-rabbit IgG | Goat polyclonal / Alexa488 | 500x | Life Technologies | A11034 |
| Anti-mouse IgG | Goat polyclonal / Alexa555 | 500x | Life Technologies | A21424 |
| Anti-rabbit IgG | Donkey polyclonal/Alexa647 | 500x | Life Technologies | A31573 |

#### Immunofluorescence of cultured PGCLC-derived cells

Immunofluorescence analysis of cultured PGCLC-derived cells was performed as described in (Nagaoka *et al*, 2020). Briefly, PGCLCs were cultured on m220 feeder cells seeded on a 0.1% gelatin-coated plate used specifically for imaging (Nunc, 165305). PGCLC-derived cells were fixed at c5, c7 or c9 with 4% PFA (Electron Microscopy Science, 15713) in PBS at room temperature for 30 min, followed by three washes in PBS. Fixed cells were blocked in PBS containing 10% NGS, 3% BSA, and 0.2% Triton X-100 for 1 hr, then incubated with the primary antibodies diluted in a 1:1 solution of blocking buffer to PBS with 0.2% Tween (PBST) at room temperature overnight. After three washes in PBST, cells were incubated with the secondary antibodies and DAPI at room temperature for 2 hr and washed three times in PBST. Finally, the well was filled with VECTASHIELD without DAPI (Vector laboratories, H1000). Immunostained samples were observed with a Leica SP8 confocal microscope.

#### Meiotic cell spreads

Cultured PGCLC-derived cells were harvested by TrypLE Express at 37°C for 5 min, quenched with 1:1 TrypLE wash buffer (DMEM/F12 containing 0.1% BSA fraction V, 30 mM HEPES), filtered through a 70 µM strainer and centrifuged at 1200 rpm for 5 min. Cell pellets were dislodged by tapping and washed once in PBS. Cells were then treated with a hypotonic solution (30 mM Tris-HCl, 50 mM sucrose (Sigma, S0389), 17 mM trisodium citrate, 5 mM ethylenediaminetetraacetic acid (EDTA), 2.5 mM dithiothreitol (DTT) (Sigma, D0632), 0.5 mM phenylmethylsulfonylfluoride (PMSF) (Sigma, P7626), pH 8.2-8.4 at room temperature for 20 min. Cells were spun down 3 min at 1200 rpm, resuspended in 100 mM sucrose and the cell suspension distributed onto slides (Matsunami, S9901) covered with 1% PFA in H_2_O (Electron Microscopy Science, 15713) with 0.2% Triton X-100 (pH 9.2-9.4). The slides were incubated at room temperature overnight in a humidified chamber. Finally, the slides were air-dried and washed with 0.5% Kodak Photo-Flo 200 (Kodak, B00K335F6S) for 2 min at room temperature. The spread slides were blocked in PBS containing 10% NGS, 1% BSA for 1 hr, then incubated with the primary antibodies diluted in a 1:1 solution of blocking buffer to PBS with 0.2% Tween (PBST) at room temperature overnight. After three washes in PBST, cells were incubated with the secondary antibodies and DAPI at room temperature for 2 hr, washed three times in PBST and mounted in VECTASHIELD mounting medium with DAPI (Vector Laboratories, h1200). Immunostained cells were observed under a Leica SP8 confocal microscope.

#### Immunofluorescence of IVDi tissues

Day 11 IVDi tissues were treated while still attached to the transwell member as follow: culture medium was carefully removed from the transwell and the whole membrane was fixed in 4% PFA (Electron Microscopy Science, 15713) in PBS for 30 min at room temperature, washed twice with PBS and blocked overnight at room temperature in 10% NGS, 1% BSA, 0.2% Triton X-100. Primary antibodies were diluted in a 1:1 solution of blocking buffer to PBS with 0.2% Tween (PBST) and incubated overnight. After 3 washes with PBST, secondary antibodies and DAPI diluted as the primary, were incubated an additional overnight, washed thrice and the whole membrane mounted on VECTASHIELD with DAPI (Vector Laboratories, H1200). Immunostained tissues were observed under a Leica SP8 confocal microscope.

#### RNA-fluorescent *in situ* hybridization and immunofluorescence

Cells were fixed with 3% paraformaldehyde PFA (Electron Microscopy Science, 15713) for 10 min with 2 mM Ribonucleoside-Vanadyl Complex RVC (New England Biolabs, S1402S) at room temperature and then permeabilized for 5 min on ice in 0.5% Triton-X with 2mM RVC. Cells were then blocked in 3% BSA/PBS with 2mM RVC for 1h at room temperature, incubated with primary antibodies diluted in blocking solution with 2mM RVC overnight at 4°C. The secondary antibodies were diluted in blocking buffer and incubated 1h at room temperature. Cells were then again fixed in 3% PFA for 10 min at room temperature. Strand-specific RNA FISH was performed with fluorescently labeled oligonucleotides (IDT) as described previously (Del Rosario *et al*, 2017). Briefly, probe mix was prepared by mixing 10 ng/ml equimolar amounts of Cy5 labeled Xist probes BD384-Xist-Cy5-3’ (5’-ATG ACT CTG GAA GTC AGT ATG GAG /3Cy5Sp/ -3’), BD417-5’Cy5-Xist-Cy5-3’ (5’-/5Cy5/ATG GGC ACT GCA TTT TAG CAA TA /3Cy5Sp/ -3’), 0.5 µg/µL yeast t-RNA (Life Technologies, 15401029) and 20 mM RVC. Probe mix was pre annealed at 80°C for 10 min followed by 30 min at 37°C and hybridized in 25% formamide, 10% dextran sulfate, 2xSSC pH 7 at room temperature overnight. Slides were then washed in 25% formamide 2xSSC pH 7 at room temperature, followed by washes in 2xSSC pH 7 and then mounted with Vectashield (Vector Laboratories, H1200). Images were acquired using a Zeiss Cell Observer.

#### RNA extraction, cDNA synthesis and qPCR analysis

Total RNA was isolated from ESCs, EpiLCs, and PGCLCs (two biological replicates each, corresponding to two different clones, with further two technical replicates each) using phenol-chloroform extraction (Sigma Aldrich, P2069) followed by ethanol precipitation and quantified by Nanodrop. cDNA was produced with a High-Capacity RNA-to-cDNA Kit (Thermo Fisher Scientific, 4387406) and was used for qRT-PCR analysis in triplicate reactions with Power SYBR Green PCR Master Mix (Thermo Fisher Scientific, 4367659). The gene expression levels are presented as ΔΔCt normalized with the mean Ct values of one housekeeping gene, Arbp, in a normalization sample (ESCs). The primer sequences used in this study are listed in Table 2.

#### Bulk RNA-Seq analysis

RNA libraries were prepared using the TruSeq Stranded Total RNA Library Preparation Kit (Illumina, 20020596) followed by 125 bp paired-end sequencing on an Illumina HiSeq 2500.

#### Allele-specific Analysis

FastQ files that passed quality control were aligned to the mm10 reference genome containing CAST/EiJ and 129S1/SvImJ SNPs positions masked. The positions of all 36 mouse strains SNPs were downloaded from ftp://ftp-mouse.sanger.ac.uk/REL-1505-SNPs_Indels/mgp.v5.merged.snps_all.dbSNP142.vcf.gz.tbi. From here, we generated a VCF file containing only the SNPs information for the strains of interest, CAST/EiJ and 29S1/SvImJ. Reads with >= 1 SNPs were retained and aligned using STAR (Dobin *et al*, 2013) implementing the WASP method (van de Geijn *et al*, 2015) for filtering of allele-specific alignments.

The generated bam files were used for counting reads using the HTseq tool (v0.6.1) (Anders *et al*, 2015). All of the steps above were performed using a customized Nextflow pipeline (Di Tommaso *et al*, 2017). We obtained between 50×10^6^ and 75×10^6^ reads per replicate. Coherence between samples, time points and replicates was verified by principal component analysis (PCA). Batch effects in principal component analysis (PCA) for comparison to *in vivo* samples were corrected using the R package limma (Ritchie *et al*, 2015).

Differential expression analysis was performed using the R package DESeq2 (v1.16) (Love *et al*, 2014). Briefly, differentially expressed genes were called by comparing XGFP+ PGCLCs and XGFP-PGCLCs or XGFP+ PGCLCs to ESCs. The DESeqDataSet (dds) was generated considering the dataset in its entirety while the DEseq analysis was conducted on dataset filtered as follows: Read counts were normalized by library size using “estimateSizeFactors*”*, were filtered for having a mean across the samples >10 (a more stringent cut off than the sum across the samples >10) and poorly annotated genes on chromosomal patches were removed. The resulting 16289 genes were kept for downstream analysis. Log2 fold change was shrinked using the “normal” parameter.

Gene Ontology enrichment analysis performed on top and bottom differentially expressed genes defined as FDR < 0.001 e log2 fold change > |1| using the Gorilla. Over-represented categories were simplified using Revigo (http://revigo.irb.hr/) using a similarity of 0.4 as threshold. As background, all identified genes were used.

#### Single-cell RNA-seq analysis

Full-length single-cell RNA-seq libraries were prepared using the SMART-Seq v5 Ultra Low Input RNA (SMARTer) Kit for Sequencing (Takara Bio). All reactions were downscaled to one-quarter of the original protocol and performed following thermal cycling manufacturer’s conditions. Cells were sorted into 96-well plates containing 2.5 µl of the Reaction buffer (1× Lysis Buffer, RNase Inhibitor 1 U/µl). Reverse transcription was performed using 2.5 µl of the RT MasterMix (SMART-Seq v5 Ultra Low Input RNA Kit for Sequencing, Takara Bio). cDNA was amplified using 8 µl of the PCR MasterMix (SMART-Seq v5 Ultra Low Input RNA Kit for Sequencing, Takara Bio) with 25 cycles of amplification. Following purification with Agencourt Ampure XP beads (Beckmann Coulter), product size distribution and quantity were assessed on a Bioanalyzer using a High Sensitivity DNA Kit (Agilent Technologies). A total of 140 pg of the amplified cDNA was fragmented using Nextera XT (Illumina) and amplified with double indexed Nextera PCR primers (IDT). Products of each well of the 96-well plate were pooled and purified twice with Agencourt Ampure XP beads (Beckmann Coulter). Final libraries were quantified and checked for fragment size distribution using a Bioanalyzer High Sensitivity DNA Kit (Agilent Technologies). Pooled sequencing of Nextera libraries was carried out using a HiSeq4000 (Illumina) obtaining between 0.5×10^6^ to 1.5×10^6^ reads per cell. Sequencing was carried out as paired-end (PE75) reads with library indexes corresponding to cell barcodes.

Allele-specific alignment was done as described for bulk RNA-seq analysis using STAR and WASP. Data Processing and visualization was performed using the R package Seurat (v4.0) (Stuart *et al*, 2019). Low-quality cells with less than 4,000 identified genes, less than 10,000 RNA molecules or more than 5% mitochondrial reads were removed. Data was log normalized and the top 2,000 highly variable features were selected for downstream analysis. The expression matrix was then scaled and linear dimensional reduction was performed. To ensure that our analysis wouldn’t be confounded by *in vitro* differentiation artefacts, we focussed our analysis on germ cells by subsetting for cells with a normalized and scaled *Dazl* expression greater than 1 (60 out of 460 sorted germ cells did not pass this criterium). Moreover, only cells that passed our allelic expression QC (explained below) were retained. Clusters were subsequently identified using “FindClusters” at a resolution of 0.8 on the first 20 principal components and visualised as UMAP projections using “RunUMAP”. Clusters were annotated based on marker gene expression. Processing of allelic data was performed for all cells that passed the Seurat QC. Cells that passed the following criteria were considered for downstream analysis: More than 3,500 total allelic reads (sum of mus and cas), a minimum of 25 allelically expressed genes as well as a minimum of 3% of total allelic reads from either genotype. Moreover, a gene was considered informative if the sum of its allelic reads was higher than 10 and if it was expressed in at least 25% of cells. This resulted in 379 cells that passed all our quality control steps.

#### RNA velocity analysis

Non-allele specific RNA velocity analysis was performed as follows: Briefly, *loom* files only of *Dazl* positive cells were generated from the non-allelic specific BAM files from STAR using *velocyto run-smartseq2* version v0.17.17 using the default parameters, mouse genome assembly mm10, and the UCSC repeat genome masked regions using custom made scripts.

Subsequently the *loom* files were imported into Python version 3.7 and processed using scVelo v0.2.3 (Bergen *et al*, 2020). The metadata, the clusters and the UMAP dimensionality reduction coordinates from Seurat were imported then the single-cell data were filtered and normalized with a minimum of 20 counts and 2000 top genes. The moments for velocity estimations were computed with 20 principal components and 30 neighbours. The genes’ full splicing kinetics were recovered before estimating the velocities using the *dynamical model*. The RNA-velocity was visualised using *velocity_embedding_stream* colour coding cells by their Seurat cluster.

#### Integration with *in vivo* datasets

Single-cell data of *in vivo* female germ cells was obtained from GEO GSE130212 (Zhao *et al*, 2020). Data Processing and visualization was performed using the R package Seurat (v4.0) (Stuart *et al*, 2019). Low-quality cells with less than 2,000 identified genes, less than 2,000 RNA molecules or more than 5% mitochondrial reads were removed. Data was log normalized, the top 2,000 highly variable features were selected for downstream analysis and the expression matrix then scaled. Normalized and scaled *in vivo* and *in vitro* data from this study were merged by canonical correlation analysis (CCA) using the Seurat function RunCCA. UMAP was then performed using CCA.

## Data availability

Raw and pre-processed data generated will be available upon peer-reviewed publication at GEO under accession: GSE169201 (https://www.ncbi.nlm.nih.gov/geo/query/acc.cgi?acc=GSE169201).

## Acknowledgements

We are grateful to present and previous B.P. lab members for discussions and input, specifically to P. Audergon and A. Tarruell for their initial contributions in the experimental design setup. We thank L. Velten for advice on single-cell RNA-seq analysis; F. Nakaki for input on integration of transcription factor vectors and H. Ohta for help in establishing the m220 expansion system. We furthermore acknowledge critical technical support by the CRG core facilities including the CRG Genomics Unit; the CRG/UPF FACS Unit; the Bioinformatics Unit; the Advanced Light Microscopy Unit and the PRBB Animal Facility. This work was supported by the Spanish Ministry of Science, Innovation and Universities (BFU2014-55275-P, BFU2017-88407-P to B.P.), the Agencia Estatal de Investigación (AEI) (EUR2019-103817 to B.P.), the AXA Research Fund (to B.P.) and the Agencia de Gestio d’Ajuts Universitaris i de Recerca (AGAUR, 2017 SGR 346 to B.P.). We would like to thank the Spanish Ministry of Economy, Industry and Competitiveness (MEIC) to the EMBL partnership and to the “Centro de Excelencia Severo Ochoa”. We also acknowledge support of the CERCA Programme of the Generalitat de Catalunya. N.A. is supported by an EMBO postdoctoral fellowship (LTF 695-2019). J.S. and M.B. were supported by La Caixa International PhD Fellowships and J.S. by a travel grant from the Company of Biologists (Development Journal).

## Author contributions

B.P., J.S. and M.B. conceived the study and wrote the manuscript with input from T.M.. J.S. and T.M. performed experiments. M.B. and J.S. established XRep cell line. J.S., M.B., N.A. and L.C. performed bioinformatic analyses. J.S., M.B. and N.A. integrated and visualized data. P.L. and H.H. prepared the single-cell sequencing libraries. N.H., K.H., Y.N., S.N. and M.S. provided constructs, mouse gonadal somatic and m220 feeder cells, and input on experimental set-up and manuscript writing. B.P. acquired funding and supervised the research.

